# *INKILN* is a novel long noncoding RNA promoting vascular smooth muscle inflammation via scaffolding MKL1 and USP10

**DOI:** 10.1101/2023.01.07.522948

**Authors:** Wei Zhang, Jinjing Zhao, Lin Deng, Nestor Ishimwe, Jessica Pauli, Wen Wu, Shengshuai Shan, Wolfgang Kempf, Margaret D Ballantyne, David Kim, Qing Lyu, Matthew Bennett, Julie Rodor, Adam W. Turner, Yao Wei Lu, Ping Gao, Mihyun Choi, Ganesh Warthi, Ha Won Kim, Margarida M Barroso, William B. Bryant, Clint L. Miller, Neal L. Weintraub, Lars Maegdefessel, Joseph M. Miano, Andrew H Baker, Xiaochun Long

## Abstract

**Background:** Activation of vascular smooth muscle cells (VSMCs) inflammation is vital to initiate vascular disease. However, the role of human-specific long noncoding RNAs (lncRNAs) in VSMC inflammation is poorly understood.

**Methods:** Bulk RNA-seq in differentiated human VSMCs revealed a novel human-specific lncRNA called <u>IN</u>flammatory M<u>K</u>L1 <u>I</u>nteracting <u>L</u>ong <u>N</u>oncoding RNA (*INKILN*). *INKILN* expression was assessed in multiple in vitro and ex vivo models of VSMC phenotypic modulation and human atherosclerosis and abdominal aortic aneurysm (AAA) samples. The transcriptional regulation of *INKILN* was determined through luciferase reporter system and chromatin immunoprecipitation assay. Both loss- and gain-of-function approaches and multiple RNA-protein and protein-protein interaction assays were utilized to uncover the role of *INKILN* in VSMC proinflammatory gene program and underlying mechanisms. Bacterial Artificial Chromosome (BAC) transgenic (Tg) mice were utilized to study *INKLIN* expression and function in ligation injury-induced neointimal formation.

**Results:** *INKILN* expression is downregulated in contractile VSMCs and induced by human atherosclerosis and abdominal aortic aneurysm. *INKILN* is transcriptionally activated by the p65 pathway, partially through a predicted NF-κB site within its proximal promoter. *INKILN* activates the proinflammatory gene expression in cultured human VSMCs and ex vivo cultured vessels. Mechanistically, *INKILN* physically interacts with and stabilizes MKL1, a key activator of VSMC inflammation through the p65/NF-κB pathway. *INKILN* depletion blocks ILIβ-induced nuclear localization of both p65 and MKL1. Knockdown of *INKILN* abolishes the physical interaction between p65 and MKL1, and the luciferase activity of an NF-κB reporter. Further, *INKILN* knockdown enhances MKL1 ubiquitination, likely through the reduced physical interaction with the deubiquitinating enzyme, USP10. *INKILN* is induced in injured carotid arteries and exacerbates ligation injury-induced neointimal formation in BAC Tg mice.

**Conclusions:** These findings elucidate an important pathway of VSMC inflammation involving an *INKILN*/MKL1/USP10 regulatory axis. Human BAC Tg mice offer a novel and physiologically relevant approach for investigating human-specific lncRNAs under vascular disease conditions.

## Introduction

Vascular homeostasis is maintained by the interplay of signaling pathways between resident and circulating cells. Humoral, physical, or mechanical perturbations to the vessel wall trigger inflammation.^1^ Sustained and over-activated vascular inflammation leads to vascular cell maladaptation and pathological vascular remodeling.^1^ Excessive vascular inflammation underlies virtually all pathological events in the vasculature, including neointimal formation, lipid accumulation, plaque destabilization, aortic rupture, and thrombosis.^2^ Targeting vascular inflammation is considered a promising strategy to combat different vascular disorders as evidenced by numerous preclinical animal trials as well as the CANTOS Trial.^3–7^ Despite these efforts, successful implementation of anti-inflammatory strategies for vascular diseases remains disappointing.^8, 9^ This is likely due to the challenges in selectively targeting vascular inflammation among the highly complex network of systemic inflammatory pathways. As such, a better understanding of the molecular underpinnings of vascular inflammation, particularly as to how diverse coding and noncoding genes intertwine to govern this process, is essential to develop effective anti-inflammatory based therapeutics for vascular disease.

The human genome undergoes pervasive transcription of long non-coding RNAs (lncRNAs), defined as processed transcripts of length ≥ 200 nucleotides with no protein coding potential.^10, 11^ Unlike mRNAs, which are highly conserved across mammalian species, the majority of lncRNAs are human-specific, precluding loss-of-function studies in rodent models. Recent studies identified numerous human-specific lncRNA associated with complex cardiometabolic traits.^11, 12^ Characterization of these potentially important human non-conserved lncRNAs in cardiovascular pathophysiology remains challenging due to the lack of *in vivo* models. Engineering mice with human Bacterial Artificial Chromosomes (BACs) carrying human sequences, especially human lncRNA gene loci, represents a potentially effective approach to this endeavor.^13, 14^ However, this approach has yet to be harnessed for *in vivo* investigation of lncRNAs under vascular disease contexts.

A number of lncRNAs have been documented as key regulators in various biological processes and human diseases.^15^ The actions of lncRNAs depend on their cellular localization, which confers the physical accessibility to their interactive partners for function. Nuclear lncRNAs can modulate gene expression by associating with DNA, transcription factors, and epigenetic modifiers while cytosolic lncRNAs partner with diverse factors to influence protein translation, RNA or protein stability, and protein activity.^16^ Recent efforts have revealed the important roles of lncRNAs in vascular pathophysiology, such as VSMC differentiation, angiogenesis, oxidative stress, senescence, endothelial permeability, and, more recently, endothelial to mesenchymal transition.^17–22^ Several lncRNAs have been reported to regulate vascular inflammation. For example, the human specific *lncRNA-CCL2* positively regulates its neighboring protein coding gene, *CCL2*.^23^ LncRNA *VINAS* promotes atherosclerosis via activating NF-κB and MAPK signaling pathways.^24^ Hematopoietic *MALAT1* inhibits vascular inflammation through sponging microRNA miR-503.^25^ While these studies have been conducted in endothelial cells and macrophages, lncRNA function in VSMC inflammation remains mostly unknown. Elucidating inflammatory lncRNAs in VSMC phenotype transition is of critical importance given the established role of VSMCs in such inflammatory vascular diseases as atherosclerosis and aneurysm.^26^

The widely expressed Myocardin related transcription factor A (MRTFA, MKL1) is a multifaceted transcription factor, initially recognized as a cofactor of SRF to facilitate CArG-dependent gene transcription of the VSMC contractile gene program.^27, 28^ In contrast to MYOCD and MRTFB (MKL2), whose function is to establish and maintain VSMC differentiation,^29^ MKL1 expression is robustly induced by and contributes to diverse vascular pathologies, including neointimal formation, atherosclerosis, hypertension, aortic dissection, and aneurysm.^30–34^ The pathological role of MKL1 in the vasculature is mediated by disparate gene programs, including extracellular matrix, oxidative stress, and vascular inflammation. ^33, 34^ The proinflammatory action of MKL1 has been attributed to its crosstalk with p65/NF-κB regulatory axis, either through physical interaction with p65 to transactivate proinflammatory genes, or complex with epigenetic modifiers to confer an active chromatin state around proinflammatory.^35, 36^ As such, it is of particular importance to understand how MKL1 is pathologically induced. However, the mechanism underlying MKL1 expression in vascular disease contexts, particularly at the protein level, is virtually unknown.^37^

In the present study, through an unbiased RNA-seq study, we report on a novel human-specific lncRNA, called <u>IN</u>flammatory M<u>K</u>L1 <u>I</u>nteracting <u>L</u>ong <u>N</u>oncoding RNA or *INKILN.* We show that *INKILN* is positively associated with and activates proinflammatory VSMC phenotype via a MKL1/p65 pathway. This involves *INKILN* physical interaction with MKL1 and a deubiquitinase, USP10, to prevent ubiquitin-dependent MKL1 degradation. Using an innovative human BAC *INKILN* transgenic mouse model, we demonstrate that *INKILN* is induced by and contributes to neointimal formation in a carotid artery ligation model. In addition to revealing a new lncRNA and molecular mechanism underlying the proinflammatory VSMC phenotype and vascular pathology, we present an innovative BAC transgenic approach to study the *in vivo* regulation and function of human-specific lncRNAs in models of human disease.

## Methods and Material

### Cell culture, human saphenous vein ex vivo culture, and siRNA knockdown

Primary human aortic, coronary artery, pulmonary artery SMCs, saphenous vein SMCs (HASMCs, HCASMCs, PASMCs, HSVSMCs) and other human cells, such as human rhabdomyosarcoma (RD) cells, HITB5, and SMC-like myofibroblasts (BR5) were prepared and maintained as previously described.^17, 38^ Discarded human saphenous vein (HSV) tissues were obtained from patients undergoing coronary artery bypass grafting (CABG) procedure. Sample collection and the related experiments were exempted from human subject restrictions by institutional Review Board (IRB) at Augusta University. Freshly collected HSV samples were immediately processed for either ex vivo culture or primary smooth muscle cell dispersion. For ex vivo culture, HSV samples were placed in sterile PBS buffer, longitudinally opened, and transversely cut into 5 mm segments. Each segment was pinned onto a sterile surgical mesh with luminal surface facing up in a well of 12-well plate. Vein segments were cultured for 2 weeks before RNA isolation for qRT-PCR analysis of proinflammatory gene expression.

Duplex siRNA (DsiRNA) knockdown in ex vivo HSV segments was carried out as reported recently.^39^ Briefly, Duplex siRNA targeting *INKILN* or a negative control Duplex siRNA were reconstituted by PBS at a final concentration of 25 μM. Vein segments were soaked in siRNA solution for 30 minutes, rinsed with PBS, and cultured in RPMI1640 supplemented with 10% FBS and 1% MycoZap Plus-PR (LONZA, VZA-2021) for 3 days before RNA extraction and qRT-PCR analysis. siRNA knockdown in human primary cells was conducted as described elsewhere.^17^ DsiRNA targeting *INKILN* or negative control DsiRNA were purchased from Integrated DNA Technologies (IDT). ON-TARGETplus SMARTpool siRNAs targeting MKL1 (Cat. # L-015434-00-0020), USP10 (Cat. # L-006062-00-0020), and scramble siRNA control (D-001810-10-20) were all purchased from Dharmacon. FANA modified antisense oligonucleotides were purchased from AUM Biotech. Working concentration for both DsiRNAs and Dharmacon SMARTpool siRNA in cultured cells was 25 nM, and FANA ASO was 100 nM. Lipofectamine 2000 (Invitrogen) was used as the transfection reagent per the manufacturer’s instruction. RNA or protein was isolated at 48-72 hours post transfection. Treatment of the human cells with different proinflammatory stimuli was performed as described previously.^38^

### Human vascular disease samples

Human abdominal aortic aneurysm (AAA) and atherosclerotic carotid endarterectomy (CEA) samples were provided by the Munich Vascular Biobank. The AAA samples were collected from patients undergoing elective open AAA repair and the control aortas were from kidney transplant surgery organ donors. The detailed information of AAA samples is described elsewhere.^34^ The experiments were conducted in accordance with the related human subject study protocol which was approved by the Institutional Review Board (IRB) at the Klinikum rechts der Isar of the Technical University Munich. Total RNA isolation and qRT-PCR were conducted per standard methods described below. The characteristics of patients for AAA were described previously ^34^ and for CEA specimens are included in **Supplementary Table 1**.

### Bulk RNA sequencing in HASMCs and data analysis

Bulk RNA-seq in HCASMCs overexpressing MYOCD and data analysis were described previously and related data were deposited in Gene Expression Omnibus (GEO) database (GSE77120).^17^ Bulk RNA-seq for *INKILN* knockdown was conducted in human aortic SMCs (HASMCs). Briefly, HASMCs were treated with two independent DsiRNAs targeting *INKILN* or a negative control RNA using Lipofectamine 2000 at a concentration of 25 nM for 36 hours. Cells were then starved overnight followed by IL1β (4ng/ml) or vehicle control treatment for 24 hours before total RNA extraction. RNA was submitted to Genomics Research Center at University of Rochester Medical Center for deep sequencing. Polyadenylated RNA was selected for deep sequencing at a depth of 20 million reads per replicate using Illumina HiSeq 2500 system (Illumina, San Diego, CA, USA). Detailed information on library construction and sequence depth was described previously^17^. Raw reads generated from the Illumina base calls were demultiplexed by bcl2fastq version 1.8.4. We used FastP version 0.20.0 to perform quality filtering and adapter removal. Processed reads were mapped to the Homo sapiens reference genome (hg38 + Gencode-31 Annotation) by using STAR_2.7.0f. DESeq2-1.22.1 was utilized to conduct differential expression analysis with a p-value threshold of 0.05 within R version 3.5.1 (https://www.R-project.org/). The PCA plot was created through the pcaExplorer for measuring sample expression variance. Heatmaps were generated using the pheatmap package. Gene ontology analyses were performed using the EnrichR package. The detailed RNA-seq information of this assay is available in GSE158219 deposited in NIH GEO database.

### Coronary artery single nucleus (sn) ATAC-seq data analysis

SnATAC-seq was performed on coronary artery samples from 41 patients as described previously using the commercial 10x Genomics Single Cell ATAC platform.^40^ Coronary artery segments were obtained from explanted hearts from transplant recipients and hearts that were rejected for orthotopic heart transplantation (Stanford University School of Medicine). All samples were de-identified, and subjects provided consent through Stanford University IRB-approved protocols for research study participation. Proximal coronary artery segments from either the left anterior descending artery, left circumflex artery, or right coronary artery were placed in cardioplegic solution before dissection. After dissection, samples were flash frozen in liquid nitrogen and stored at −80°C. For the snATAC-seq experiments, all coronary samples were broken into small fragments using a chilled mortar and pestle with dry ice and liquid nitrogen. Nuclei were then isolated using a Iodixanol/sucrose gradient and density gradient centrifugation based on the Omni ATAC-seq protocol.^41^ After isolation, nuclei were first transposed with Tn5 transposase in bulk, followed by capture of gel beads in emulsions (GEMs) using the 10x Genomics Chromium Controller instrument. Single nucleus ATAC-seq libraries were prepared using the 10x Genomics Chromium Single Cell ATAC Kit (version 1 or 1.1 (Next GEM)). Barcoded libraries were sequenced on an Illumina NovaSeq 6000. Coronary artery snATAC-seq data was preprocessed using the 10x Genomics Cell Ranger ATAC pipeline (version 1.2.0) and reads mapped to the hg38 reference genome. Downstream snATAC-seq analyses were performed using the ArchR software package (version 1.0.1) to retain high quality/informative nuclei with transcription start site (TSS) enrichment >= 7 and >=10000 unique fragments.^42^ Chromatin accessibility profiles were linked to gene expression by integrating snATAC-seq data with human coronary artery scRNA-seq data from 4 individuals to annotate cell types.^43^ Pseudo-bulk data for each labeled cell type were visualized on the UCSC genome browser. For disease-specific analyses, the chromatin accessibility profiles from donors at different disease stages were aggregated based on three defined categories as previously described.^40^ Information on sample categorization was described in the related figure legend. The genomic region for *INKILN* was mapped to coronary artery snATAC profiles in UCSC using BLAT and the hg38 genome build.

### RNA isolation and quantitative reverse transcriptase-PCR (qRT-PCR)

HSV segments were mixed with Trizol per Precellys Lysing Kit (VWR Scientific, Radnor, PA) and homogenized by a Minilys homogenizer (Bertin Technologies, Rockville, MD). We used miRNeasy mini kit (Qiagen) to extract total RNA from homogenized vessels and cultured cells. After concentration measurement by a Nanodrop 2000 spectrophotometer (Thermo Fisher Scientific), 200-500 nanograms (ng) of total RNA were used to synthesize cDNA following an iScript cDNA kit (Bio-Rad). Universal SYBR Green Supermix (Bio-Rad) and CFX386 Touch™Real-Time PCR Detection System (Bio-Rad) were utilized to conduct qRT-PCR. Sequences of primers for this assay are listed in **Supplementary Table 2**.

### Western blot analysis

Protein was extracted from cultured primary human VSMCs after different treatment or mouse carotid arteries by using ice-cold lysis buffer (Cat. # 9803, Cell Signaling Technology) supplemented with a protease inhibitor cocktail (Sigma). Equal amounts of protein samples were resolved on SDS-PAGE gel. Western blot was performed as described previously.^44^ Information on antibodies for western blot is provided in **Supplementary Table 3**.

### In vitro RNA pull down

*INKILN* V1 full length and its antisense were PCR cloned into pcDNA3.1 expression plasmids. Linearized plasmid DNA (3 ug) was utilized as template for synthesis of biotinylated RNA using AmpliScribe T7-Flash Biotin-RNA transcription Kit (Epicentre/Lucigen). The freshly synthesized biotinylated RNA was heated to 90°C for 2 minutes, chilled on ice for 2 minutes, mixed with RNA structure buffer (10 mM Tris pH 7, 0.1 M KCl, 10 mM MgCl_2_), and then shifted to room temperature (RT) for 20 minutes to allow proper secondary structure formation. Freshly collected HCASMCs pellet was re-suspended in 1 ml RIP buffer (150 mM KCl, 25 mM Tris pH 7.4, 0.5 mM DTT, 0.5% NP40, 1 mM PMSF and protease Inhibitor (Roche, Cat. #11836153001) followed by 10 minutes centrifugation at 13,000 RPM for 10 minutes. Supernatant was collected to incubate with the folded RNA at RT for 1 hour and then immobilized by a streptavidin agarose beads (Invitrogen) at RT for another 1 hour. Beads were washed briefly for five times and boiled in SDS buffer followed by SDS page electrophoresis. Silver staining was conducted per manufacture’s instruction (Thermo Scientific, Cat. #24612) and standard western blot analysis was performed as described above. Antibody information for this assay is listed in **Supplementary Table 3.**

### RNA immunoprecipitation

Magna RIP™ RNA-Binding Protein Immunoprecipitation (RIP) Kit (Millipore Sigma, Cat. #17-700) was used for this assay with minor modification of vendor’s manual. Primary human VSMCs or a rhabdomyosarcoma cell line (RD) were used for this assay. Briefly, 2×10^7^ growing cells were harvested and lysed in 100 μl lysis buffer for one reaction of immunoprecipitation (IP) and temporarily stored in - 80°C. Freshly thawed cell lysates were sonicated with a Bioruptor at high magnitude for 2 minutes (20 seconds on, 1 minute off) followed by a brief centrifugation. The supernatant was transferred to a new tube for precipitation of RNA. Magnetic beads were incubated with 5 μg of antibody of interest for 30 minutes at RT. The conjugated beads were then mixed with 100 μl lysate supernatant and 900 μl RIP buffer followed by an overnight incubation at 4°C. Protein-RNA complex was retrieved on a magnetic separator and incubated with Proteinase K to remove protein. RNA was extracted by the phenol/chloroform prior to standard RT-PCR assay to assess the enrichment of RNA(s) of interest. Information on the antibodies is listed in **Supplementary Table 3**.

### Immuno-RNA fluorescence in situ hybridization (FISH)

RNA FISH in cultured VSMCs was carried out according to the protocol provided by Affymetrix QuantiGene View RNA ISH Cell Assay kit with certain modification for co-immunostaining of proteins. Primary human VSMC cells were seeded on 12 mm round coverslips in tissue culture plates. Cells were fixed by 4% formaldehyde at RT, rinsed with PBS, and permeabilized by 0.5% Triton X-100/PBS containing 5mM Ribonucleoside vanadyl complexes (VRC, Sigma-Aldrich, Cat. #94740). Cells were rinsed with PBS followed by the procedures for RNA FISH as described previously.^17^ Briefly, cells were sequentially incubated with PreAmplifier, Amplifier, and Label Probe at 40°C for 30 minutes, respectively. Cells were then rinsed with wash buffer and subjected to Post-FISH fixation using freshly prepared 4% formaldehyde. Cells were finally blocked with 5% BSA at RT for 1 hour prior to immunofluorescence staining and confocal imaging as described previously.^17^ Staining of *INKILN* and ACTA2 (Abcam, Cambridge, UK) in human AAA tissue was performed using RNAscope Multiplex Fluorescent v2 Assay combined with immunofluorescence staining (ACD, Biotechne). Positive and negative control probes were used in each round. Formalin-fixed paraffin-embedded human AAA tissues were sectioned for 3 μm. Sections were blocked with hydrogen peroxide followed by heat-mediated antigen retrieval for 20 minutes and Protease Plus treatment for 30 minutes. Sections were then incubated with *INKILN* or negative control probe (sequence available upon request) at 40°C for 2 hours in an oven (Boekel Sciantific, Feasterville-Trevose, US). Signal amplification (AMP1-3) and development of HRP with fluorophore Opal 690 (Akoya Bioscience, Marlborough, US) were performed per the manufacturer’s instructions prior to ACTA2 immunostaining as previously described.^45^ Images were acquired with Olympus FLUOVIEW FV3000 confocal microscope. RNA-FISH for mouse vessels was conducted using RNAScope™ ISH Kit from ACD as recently reported.^46^ Freshly collected mouse carotids were fixed in 10% neural buffered formalin (NBF) and 5 μm paraffin sections were used. The ISH signal was amplified using AMP solution and the final color was developed through FAST RED solution. Images were acquired by LSM900 confocal microscope (Zeiss). Information on related antibodies is listed in **Supplemental Table 3**.

### Proximity ligation assay (PLA)

PLA was used to detect the physical interaction between two proteins in cultured HCASMCs per Duolink^®^ PLA Fluorescence Protocol (Sigma). Growing cells were transfected with *siINKILN* or the negative control siRNA for 3 days and then incubated sequentially with −20°C ethanol for 10 minutes, 0.2% saponin for 15 minutes, and Duolink^®^ Blocking for 1 hour. Cells were then incubated with mouse anti-MKL1 primary antibody and rabbit anti-USP10 primary antibody at 4°C overnight followed by incubation with anti-rabbit PLUS and anti-mouse MINUS 1/5 diluted PLA probes (as secondary antibodies) at 37°C for 1 hour. These secondary antibodies are coupled with oligonucleotides targeting primary antibodies. Cells were then incubated with 1/40 diluted ligase at 37°C for 30 minutes to hybridize the connector oligonucleotides so that the PLA probes, which are in close proximity, form a closed circular DNA template. Cells were then incubated with amplification buffer diluted DNA polymerase at 37°C for 100 minutes to amplify the closed circular DNA template using PLA probe as a primer. After final washes, cells were mounted with Duolink^®^ In Situ Mounting Media. Negative controls were performed by incubation with either MKL1 or USP10 primary antibody alone. PLA signals were acquired by LSM900 confocal microscope (Zeiss). Information on antibodies is listed in **Supplemental Table 3**.

### Ubiquitin assay and co-immunoprecipitation (Co-IP)

For ubiquitin assay, cells were treated with MG132 (10 μM)-containing culture medium for the last 6 hours prior to protein extraction. VSMC lysates from one 10-cm dish were used for one reaction of IP experiment. Cells were washed twice with ice cold PBS and lysed with 400 μl cell lysis buffer (Cell Signaling, Cat. # 9803) supplemented with protease inhibitor cocktails (Roche, 11836153001) and NEM (N-ethylmaleimide). Cells were incubated in lysis buffer with gentle shaking at 4°C for 1 hour. Cell lysates were centrifuged briefly and supernatant was collected for immunoprecipitation by incubating with antibodies of interest overnight at 4°C. Protein precipitates were immobilized by magnetic beads (NEB, Cat. # S1432S). Beads were then washed with lysis buffer and boiled with sample buffer (Bio-Rad, 1610747) at 95°C for 10 minutes. Samples were collected for western blot of targeting proteins. Information on the antibodies is listed in **Supplemental Table 3**.

### Lentiviral particle preparation and viral transduction

Rapid Amplification of cDNA Ends (RACE) kit (Ambion) was utilized to determine the 5’ and 3’ ends of *INKILN* _V1 and *INKILN_V2* transcripts. The full-length *INKILN_V1* transcript was cloned into pcDNA3.1 vector and confirmed by DNA sequencing. The lentivirus carrying the *INKILN_V1* transcript and vector control lentivirus were generated using pFUGW/pCMV-VSVG/pCMV-dR8.2 packaging system as described previously.^17^ Lentiviral vector pLKO.1 carrying short hairpin RNA targeting human *MKL1* was obtained from Addgene (plasmid #27161). Control vector with non-mammalian shRNA was purchased from Sigma (Cat. # SHC002). Lentiviral particles were packaged in HEK293FT cells with co-transfection of vector plasmid along with packaging plasmid pCMVΔ6.2 and viral envelope-encoding plasmid pCMV-VSVG as described previously.^17^ Virus-containing medium was collected 40 hours after transfection. After purification by centrifugation, virus was concentrated by the Millipore Ultra Centrifugal Filter Units (Fisher Scientific, Cat. # UFC901024). Lentiviral titer was measured by Lenti-X qRT-PCR Titration Kit (TaKaRa, Cat. # 631235). Primary VSMCs were seeded in 6-well plates at 50% density and transduced by lentivirus the next day. The detailed information on lentivirus transduction was described previously.^17^

### Chromatin immunoprecipitation (ChIP) assay

Myer lab’s ChIP protocol (online available) was used to prepare DNA for chromatin immunoprecipitation.^17^ Briefly, vehicle control or IL1β-induced HASM cells (2×10^7^) were cross-linked by formaldehyde, quenched with glycine (0.125M), rinsed with cold PBS for twice, and pelleted in Farnham lysis buffer (5 mM PIPES pH 8.0 / 85 mM KCl / 0.5% NP-40, 1% protease inhibitor). Cell pellet was re-suspended in Farnham lysis buffer and sheared with 27G needle syringe. Nuclear pellet was collected in RIPA buffer and sonicated by a Bioruptor to achieve chromatin fragments of length 300-1000 bp. Chromatin complexes were incubated with Dynabeads (Thermo Fisher, Cat. # 11203D) conjugated with either p65 antibody or the same amount of negative control rabbit IgG at 4°C overnight on a rotator. The precipitated protein/chromatin mixtures were retrieved in 200 μl elution buffer. After reverse crosslinking and column purification, precipitated DNA samples were subjected to quantitative PCR for assessing the enrichment of a NF-κB site predicted in the proximal *INKILN* promoter. Information on PCR primers and antibodies is included in **Supplemental Table 2** and **3**, respectively.

### *INKILN* in vitro transcription/translation assay

pcDNA carrying full length *INKILN_V1*, pcDNA vector control and a positive control vector expressing luciferase were utilized for in vitro transcription/translation assay by TNT Quick Coupled Transcription/Translation System (Promega, Cat. # L1171) combined with Transcend Biotin-Lysyl-tRNA label system (Promega, Cat. # L5061) as described elsewhere.^47^ In brief, 0.5 μg of above plasmid DNA template was mixed with TNT T7 Quick Master Mix, Methionine, and Transcend Biotin-Lysyl-tRNA and incubated at 30°C for 90 minutes for protein translation. The translated protein mixture was resolved in 16% Tricine-SDS-PAGE gel, and protein/peptides were detected by a Transcend Chemiluminescent Translation Detection system (Promega, Cat. # L5081).

### Luciferase assay

A 1.4 kb *INKILN* promoter fragment containing a predicted NF-κB site and a truncated 1.18 kb version without this site were PCR amplified from HCASMC genomic DNA and cloned into the pGL3 basic luciferase reporter. Luciferase assay was conducted as previously described.^48^ Briefly, HEK293 cells were plated in 24-well cell culture plates until 80% confluency before transfection of the above luciferase reporter plasmid and Renilla as an internal control. 24 hours after transfection, cells were starved overnight followed by TNFα (5 ng/ml) stimulation for 6 hours prior to luciferase activity measurement using a Dual Luciferase Assay Kit per the vendor’s protocol (Promega). To test the influence of *INKILN* on p65/NF-κB transactivity, HASMCs were treated with *siINKILN* or siCtrl (sicontrol) for 24 hours followed by electroporation of an NF-κB luciferase reporter and Renilla internal control. After overnight serum starvation, cells were stimulated with TNFα (5 ng/ml) for 24 hours prior to luciferase assay as above.

### BAC transgenic mouse generation and transgene characterization

The transgenic (Tg) *INKILN* mouse (C57BL/6 background) was generated by Cyagen carrying a human BAC clone (RP11-997L11) following deletion of the *CXCL8 (aka IL8)* gene locus through BAC recombineering.^13^ The transgenic line was initially genotyped by Cyagen via multiple PCR primers targeting different regions of the BAC transgene to ensure integrity. We also confirmed the deletion of *IL8* gene locus by gene expression and targeted genotyping. To more precisely characterize the transgene, CRISPR-LRS (Long Reads Sequencing) was used (i) to map the locus for the BAC containing *INKILN,* (ii) to determine transgene copy number, and (iii) to assess transgene-induced perturbations to the genome accompanying transgene integration.^49^ Long-read libraries with either one or two CRISPR RNA (crRNAs) were prepared using CAS9 Sequencing kit (SQK-CS9109) from Oxford Nanopore Technologies following manufacturer’s instructions (www.nanoporetech.com). For genomic DNA input, high molecular weight (HMW) genomic DNA was isolated with the Monarch HMW DNA Extraction Kit (T3060) following manufacturer’s instructions (www.neb.com). crRNAs were designed using CHOPCHOP with default parameters (https://chopchop.cbu.uib.no). Long-read libraries were run on R9.4.1 flow cells on the GridION Mk1 platform using guppy (v4.1.2) within MinKNOW (v20.10.3) and MinKNOW Core (v4.1.2) using the fast base-calling option for base-call model and a minimum Q-score of 7 for filtering of reads. Long-reads were mapped to reference sequences using the Long-Read Support (beta) plugin within Qiagen CLC Genomics Workbench (www.qiagen.com) that employs open-source tool minimap2. Lastly, mapped informative long-reads were manually queried against NCBI nr/nt, refseq genome databases (https://blast.ncbi.nlm.nih.gov/Blast.cgi) and UCSC genome browser with BLAT tool (https://genome.ucsc.edu) to obtain a working transgene integration map. To determine BAC transgene copy number, qPCR was performed with 50 ng of genomic DNA for input and data were normalized to internal control primers. Long-read sequence data have been submitted to NCBI SRA database (www.ncbi.nlm.nih.gov/sra) under BioProject number PRJNA873299. Sequence information on the crRNAs and primers for copy number analysis is listed in **Supplementary Table 4** and **5,** respectively.

### Complete ligation model, quantitation of neointimal formation, and immunofluorescence staining

All mouse studies were approved by the Augusta University Animal Care and Use Committee. *INKILN* Tg mice were crossbred with *Apoe^-/-^* mice (C57BL/6J background) for >6 generations. For more consistency in neointimal formation, 8-10 week old *INKILN Apoe^-/-^* Tg mice and their littermate control *Apoe^-/-^* mice were subjected to complete ligation at the bifurcation of left common carotid artery as previously described after 4% isoflurane inhalation-mediated anesthesia.^38^ Three weeks after ligation surgery, the ligated left and unligated right carotid arteries were carefully dissected and harvested after transcardiac saline perfusion. Carotid arteries were fixed with 10% NBF for 24 hours at 4°C with gentle shaking and then embedded into paraffin. Sections from 200 to 800 μm were prepared at a 5 μm thickness for Hematoxylin and Eosin stain (H&E) staining, RNA FISH, and immunofluorescence staining. De-paraffinization and H&E staining were carried out by Leica ST5010 AutoStainer following the manufacturer’s instructions. Images were taken by a Leica Microsystem. Neointimal and medial areas were analyzed by Image J as reported.^13, 38^ Quantitation of neointimal formation was carried out for proximal (Level 1), middle (Level 2), and distal regions (Level 3) covering 0-200, 200-400, and 400-600 μm distal to the ligature, respectively. Immunofluorescence staining for contractile smooth muscle marker ACTA2, common leukocyte marker CD45, macrophage marker MAC2, and proliferating cell marker Ki67 were carried out and quantified as previously described.^34, 38^ Information regarding the primary and secondary antibodies is listed in **Supplemental Table 3**.

### Statistical analysis

All experiments were repeated in at least 3 independent experiments. Statistical analyses were conducted with GraphPad Prism 7.0. Quantitative results were presented as mean ± standard error of mean (SEM). The normality and equal variance of the data were assessed by the Kolmogorov-Smirnov and Shapiro-Wilk tests. Unpaired t-test was used to analyze the difference between two groups. For comparison of multiple groups, two-way ANOVA followed by Sidak multiple comparison test was used for normally distributed variables. p<0.05 was considered statistically significant. The detailed information on statistical analysis is listed in each individual figure legend.

## Results

### *INKILN* expression correlates with VSMC phenotypic modulation and vascular disease

In an effort to uncover novel lncRNAs linked to VSMC phenotypic modulation and vascular disease, we conducted a bulk RNA-seq analysis in human coronary artery smooth muscle cells (HCASMs) overexpressing Myocardin (MYOCD), a potent activator of VSMC differentiation.^50^ Assembling the filtered reads derived from RNA sequencing to the human genome browser revealed numerous MYOCD-regulated protein coding genes (gray) and noncoding RNAs (red). Beyond a number of upregulated lncRNAs, such as *MYOSLID,* which we reported as an activator of VSMC differentiation,^17^ MYOCD also downregulates many lncRNAs. *INKILN* appeared to be one of the most abundantly expressed, and significantly downregulated lncRNAs (**Figure 1A**). To determine cell and tissue-specific expression of *INKILN,* we employed FANTOM (Functional Annotation Of the Mammalian genome) expression atlas, a meta-annotation which integrates reference genes and newly defined transcription start sites (TSSs) based upon CAGE-seq.^51^ Analysis of 173 cell types and 174 tissues in FANTOM revealed that *INKILN* was enriched in 5 different types of VSMCs, including SMC of the internal thoracic artery, SMC of the carotid artery, vascular associated SMC, aortic SMC, as well as different blood vessels which are enriched with VSMCs (**Supplemental Figure 1A**). Consistently, both semi- and quantitative RT-PCR (qRT-PCR) showed that *INKILN* was selectively expressed in multiple HCASMC isolates, the human VSMC cell line, HITB5, and SMC-like myofibroblasts (BR5) which were positive for the VSMC contractile gene, *LMOD1* (**Supplemental Figure 1B**). qRT-PCR experiments confirmed down-regulation of *INKILN* by MYOCD (**Figure 1B**). *INKILN* was also down-regulated by TGFβ1, another well-recognized activator of VSMC differentiation,^52^ and a commercial source of conditioned VSMC differentiation medium (SMD); like LMOD1, the VSMC contractile gene, *CNN1,* was upregulated under these conditions (**Figure 1C, 1D**). The medial layer of normal blood vessels is mainly comprised of contractile VSMCs. Upon ex vivo organ or primary cell culture, VSMCs undergo dedifferentiation, resulting in the downregulation of VSMC contractile genes.^53^ In contrast to the VSMC contractile gene *MYH11, INKILN* was undetectable in freshly obtained human saphenous veins (HSVs), whereas it was induced in ex vivo cultured HSV segments and primary HSVSMCs dispersed from the same vessel source (**Figure 1E, 1F**).

**Figure 1.**
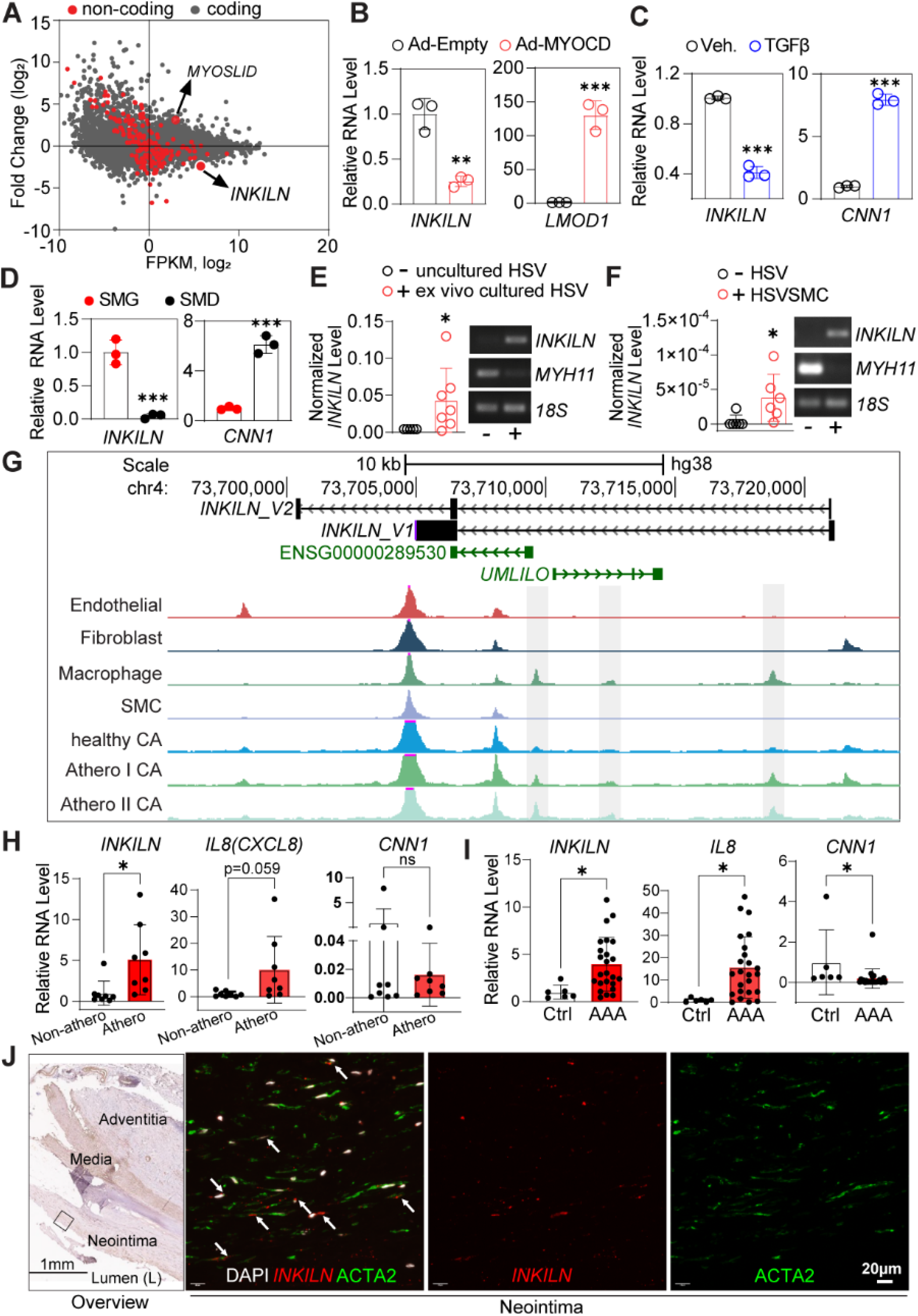
*INKILN* expression correlates with VSMC phenotypic modulation and vascular disease. **A.** RNA-seq analysis revealed numerous coding (Gray dots) and noncoding genes (red dots) regulated by MYOCD in human coronary artery smooth muscle cells (HCASMCs) (n=2). **B-D**. qRT-PCR validation of the downregulation of *INKILN* in HCASMCs transduced with Ad-MYOCD relative to Ad-empty (**B,** n=3), differentiated HCASMCs induced by either TGFβ (2 ng/ml) (**C,** n=3) or conditioned SMC differentiation medium (SMD) versus growth medium (SMG) (**D**, n=3). **E.** qRT-PCR (Left) and semi-qRT-PCR (right) analysis of the indicated genes in uncultured versus 2 weeks ex vivo cultured human saphenous vein (HSV) segments from the same patients (n=7 patients). **F.** qRT-PCR (left) and semi-qRT-PCR (right) analysis of the indicated genes in uncultured HSV versus primary cultured SMCs dispersed from fresh HSV tissues (HSVSMCs) (n=6). **G.** UCSC browser screenshot for the *INKILN* gene locus with combined single nucleus (sn) ATAC-seq libraries from healthy versus diseased coronary artery (CA) human samples (n=41). Healthy CA: patient has no evidence of atherosclerosis and samples are lesion-free; Athero I CA: patient has evidence of atherosclerosis, but samples are lesion-free; Atherosclerosis II CA: patient has evidence of atherosclerosis and sample contains lesion. **H** and **I.** qRT-PCR assessment of the indicated genes in human atherosclerotic plaque (Athero) versus non-plaque (Non-athero) regions from the same patients (**H**, n=8 patients), and abdominal aortic aneurysm (AAA) tissues (n=24 patients) relative to healthy control aortas (Control) from organ donors (**H**, n=6 donors). **J**. Representative images of the overview for the colorimetric ACTA2 (brown) immunohistochemistry staining of human AAA tissues and Immuno-RNA FISH for *INKILN* (Red) and a VSMC marker ACTA2 (Green) in the rectangle marked neointimal region (see overview) of human AAA vessels (n=5 patients). Arrows indicate specific *INKILN* signal. Unpaired two-tailed Student’s t test. *p<0.05, ** p <0.01, *** p <0.0001, ns, not significant.

Because VSMC phenotypic modulation contributes to the pathogenesis of various vascular diseases, we asked if *INKILN* expression is induced in diseased vessels. We first analyzed the combined single nucleus (sn) ATAC libraries from healthy versus diseased coronaries,^40^ and found three human atherosclerosis-associated peaks residing in intron 1 of *INKILN.* Interestingly, these peaks overlap with macrophage-specific peaks (**Figure 1G**). Further, qRT-PCR showed markedly elevated *INKILN* expression in atherosclerotic plaques compared to non-atherosclerotic regions from the same vessel source, and human abdominal aortic aneurysmal (AAA) tissues to normal aortas from healthy donors. As expected, gene expression of *CNN1* was downregulated in both atherosclerosis and AAA, which is in contrast to the proinflammatory gene *IL8***(Figure 1H, 1I)**. Immuno-RNA FISH showed a clear colocalization of *INKILN* with ACTA2 positive cells in the neointimal region of human AAA tissues, presumably representing phenotypically modulated VSMCs (**Figure. 1J**), whereas a negative control probe failed to give rise to such signal **(Supplemental Figure 1C).** Collectively, these results demonstrate that *INKILN* is a VSMC-enriched lncRNA negatively associated with the VSMC contractile phenotype and induced in vascular disease.

### *INKILN* is induced by proinflammatory stimuli through the NF-κB/p65-dependent pathway

*INKILN* is an intergenic lncRNA residing on chromosome 4, 20 kilobases upstream of *IL8*. Two splice variants of *INKILN* were found according to sequence assembly, which we refer to as V1 and V2 (**Supplemental Figure 2A**). RACE and sequencing demonstrated that the full length of *V1 and V2* is 1,750 bp and 543 bp, respectively. Because of the much lower expression of *V2* (**Supplemental Figure 2B**), we selected *V1* for overexpression experiments in this study. PhyloCSF analysis and Pfam database query support the absence of any coding potential of *INKILN* (**Supplemental Figure 2C)**. In vitro transcription/translation assay further validated the absence of coding potential in *INKILN* (**Supplemental Figure 2D**).

To determine the critical pathway(s) responsible for the induction of *INKILN* in dedifferentiated VSMCs, we analyzed our published bulk RNA-seq dataset (GSE69637) done with HSVSMCs treated with proinflammatory cytokine IL1α and growth factor PDGF, two critical stimuli driving VSMC dedifferentiation.^54^ *INKILN* was induced by IL1α but not PDGF, which was further confirmed by qRT-PCR, suggesting the proinflammatory, but not proliferative, pathway governs *INKILN* induction in VSMCs (**Figure 2A, 2B)**. Dose-dependent studies in HSVSMCs showed that high induction of *INKILN* expression was achieved by IL1α with concentrations as low as 0.01 ng/ml (**Supplemental Figure 2E**). Similar induction was seen in HCASMCs, human aortic SMCs (HASMCs), as well as pulmonary artery SMCs (PASMCs) using different proinflammatory cytokines, such as TNFα and IL1β (**Figure 2C-F, Supplemental Figure 2F).** In HASMCs, IL1β-induced *INKILN* was time-dependent with peak elevation at 12 hours following treatment. This dynamic induction paralleled its neighboring gene, *IL8* (**Figure 2G**). IL1β-induced *INKILN* was suppressed by BAY11-7082, a selective inhibitor of the NF-κB pathway (**Figure 2H**). Overexpression of IKKβ, a specific activator of the NF-κB pathway, caused a significant induction of both *INKILN* and *IL8* (**Figure 2I**), suggesting an NF-κB -dependent induction of *INKILN* expression. To determine if *INKILN* was a direct transcriptional target of the NF-κB pathway, we conducted computational analysis of the proximal promoter of *INKILN* wherein a conserved NF-κB site was predicted (**Supplemental Figure 2A**). Chromatin immunoprecipitation (ChIP) assays showed that IL1β induced p65 binding to the proximal promoter region encompassing a predicted NF-κB site in HASMCs (**Figure 2J**). Finally, TNFα significantly increased the luciferase activity of a reporter containing this NF-κB site, whereas such induction was diminished in a truncated version lacking this site (**Figure 2K**). These data support *INKILN* as a direct transcriptional target of the p65/NF-κB pathway.

**Figure 2.**
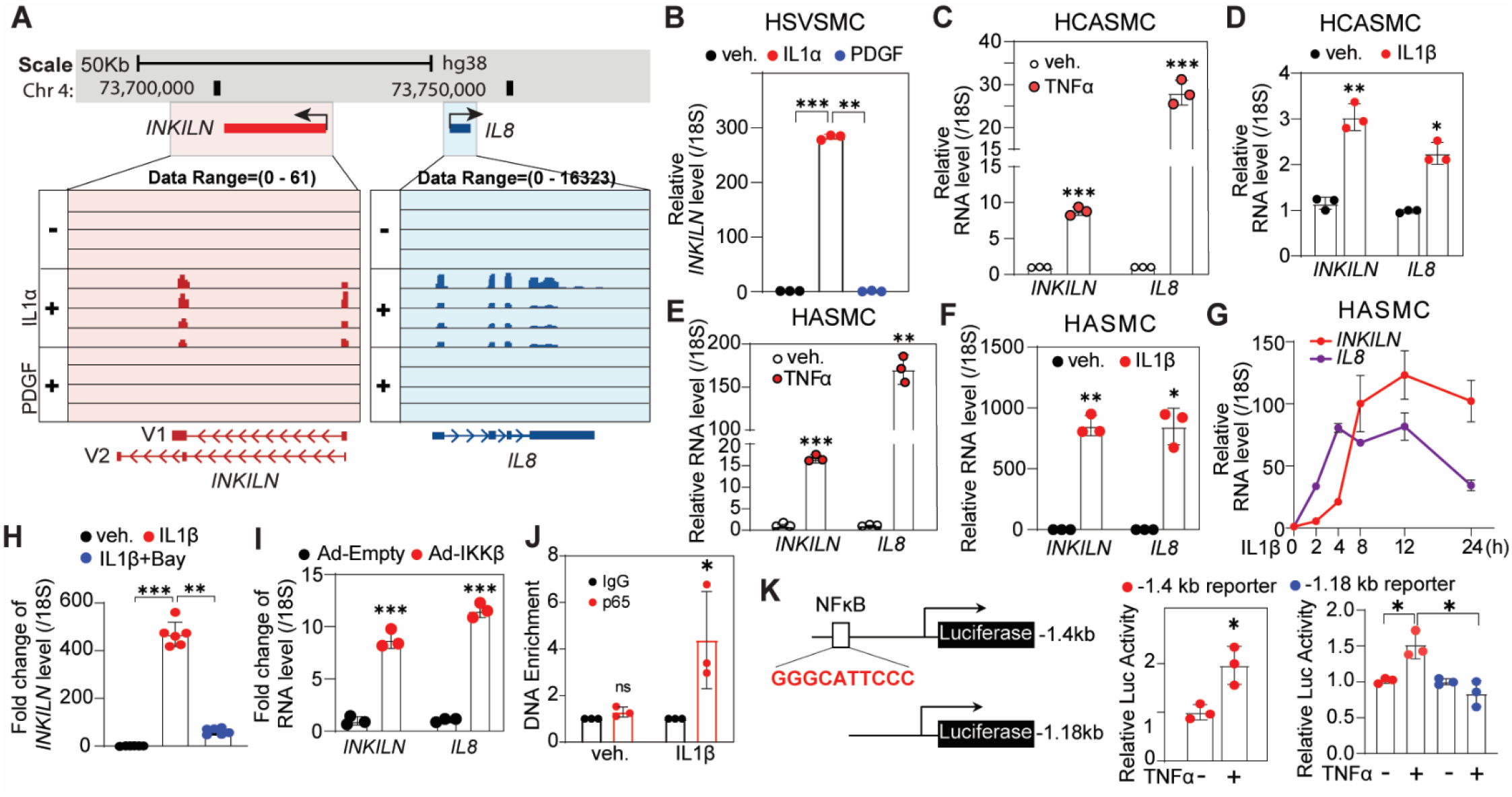
*INKILN* is induced by proinflammatory stimuli through the NF-κB/p65-dependent pathway. **A**. *INKILN* and its neighboring gene *CXCL8 (IL8)* expression in primary HSVSMCs ± IL1α and PDGF mined from the RNA-seq dataset we published.^1^ **B.** qRT-PCR analysis of *INKILN* expression in HSVSMCs induced with IL1α (10 ng/ml) and PDGF (20 ng/ml) relative to vehicle control (n=3). **C-F**. qRT-PCR assay for the indicated genes in HCASMCs ±TNFα (10 ng/ml) for 48 hours (**C**), HCASMCs ± IL1β (4 ng/ml) (**D**), human aortic SMCs (HASMCs) ± TNFα (10 ng/ml) (**E**) or IL1β (4 ng/ml) (**F)** for 24 hours (n=3). **G**. qRT-PCR for *INKILN* expression in HASMCs stimulated by IL1β (4 ng/ml) for the indicated time points (n=3). **H**. HASMCs were induced with IL1β (4 ng/ml) for 24 hours followed by treatment with BAY11-7082 (10 μM) for 24 hours before RNA extraction for qRT-PCR of the indicated genes (n=6). **I**. qRT-PCR analysis of *INKILN* in HASMCs transduced with Ad-IKKβ or vector control adenovirus (Ad-Empty) with the same dose (MOI=30) for 72 hours (n=3). **J**. Chromatin Immunoprecipitation (ChIP)-qPCR validation of p65 binding to the predicted NF-κB site within the proximal *INKILN* promoter in HASMs induced by IL1β or vehicle control for 15 minutes (n=3). **K.** Schematic of luciferase reporter of the putative −1.4 kb *INKILN* proximal promoter containing a predicted NF-κB site and the truncated - 1.18 kb reporter lacking this site, and luciferase assays for the −1.4 kb promoter of *INKILN* and the truncated reporter in HEK293 cells induced by TNFα (10 ng/ml)) for 6 hours (n=3). Unpaired two-tailed Student’s t test throughout all the panels. * p <0.05, ** p<0.01, *** p<0.0001, ns, not significant.

### *INKILN* positively regulates proinflammatory gene expression in VSMCs

The massive induction of *INKILN* by proinflammatory stimuli suggests that *INKILN* participates in the proinflammatory gene program in VSMCs. To test this hypothesis, we performed RNA-seq in HASMCs treated with two different siRNAs, individually or in combination, to *INKILN* followed by IL1β or vehicle control treatment for 24 hours. Principal Component Analysis (PCA) revealed that samples from the same condition clustered together, and differences were evident between control and *INKILN* siRNA treated groups under both vehicle and IL1β stimulated conditions (**Supplemental Figure 3A**). We next performed differential expression analysis based on an adjusted p-value ≤ 0.05 for each set of raw expression measures. A total of 548 genes were significantly differentially expressed in si*INKILN* versus siCtrl under basal conditions, with 333 of them downregulated by *siINKILN.* Notably, of 838 significantly regulated genes under the IL1β stimulated condition, 518 were downregulated (see information in **GSE158219**). To identify the pathways regulated by *INKILN* and gain insight into *INKILN* molecular functions in VSMCs, we performed Gene Ontology (GO) enrichment analysis and KEGG pathway analysis. The majority of downregulated pathways upon *siINKILN* knockdown were associated with inflammation-related biological processes, including pathways of cellular response to cytokine stimulus, cytokine-mediated signaling, chemokine-mediated signaling, and positive regulation of MAPK cascade **(Figure 3A)**. KEGG pathway analysis further identified several pathways related to inflammation and immune response, such as TNF signaling, Cytokine-cytokine receptor interaction, IL-17 signaling, and NF-kappa B signaling (**Supplemental Figure 3B)**. Numerous proinflammatory genes, including those encoding chemokines and cytokines *(CXCL1, IL8,* and *IL6),* as well as other inflammatory mediators *(NR4A2, PTGS2,* and *OLR1),*were downregulated upon *INKILN* depletion under basal and IL1β-induced conditions (**Figure 3B**). Down-regulation of the representative proinflammatory genes, such as *IL8*, *IL6*, *CCL2,* and *CXCL1* was validated by qRT-PCR in both basal and IL1β-induced HASMCs, growing HCASMCs, and TNFα-induced PASMCs (**Figure 3C, 3D**, **Supplemental Figure 3C).** These results were further validated by a separate siRNA in HASMCs **(Supplemental Figure 3D)**. To validate the siRNA results, we utilized FANA ASO, an alternative approach for gene knockdown based on RNase H-mediated RNA-degradation.^55, 56^ FANA-mediated *INKILN* knockdown resulted in a similar downregulation of proinflammatory genes in both growing HASMCs and HCASMCs, though to a lesser degree compared with siRNA, likely due to lower knockdown efficiency (**Figure 3E, 3F**). In line with the results from loss of function study, lentivirus overexpressing *INKILN* (Lenti-*INKILN)* induced *IL6*, *IL8*, *CXCL1, and CXCL5* gene expression in HASMCs (**Figure 3G**). To test if *INKILN* functions similarly in human vasculature, we utilized a well-recognized organ culture model to recapitulate pathological vein remodeling in HSV bypass grafts.^39^ Robust induction of most proinflammatory genes, including *INKILN, IL8, IL6,* and *CXCL5* was observed in HSV segments cultured for 3 days **(Supplemental Figure 3E).** Notably, siRNA was efficiently delivered to the vasculature, as evidenced by 70%*INKILN* knockdown in the cultured segments, which led to a significant reduction of *IL8, IL6,* and *CXCL5* gene expression (**Figure 3H**). Taken together, these results demonstrate that *INKILN* is a novel activator of the proinflammatory gene program in cultured VSMCs and ex vivo cultured vessels.

**Figure 3.**
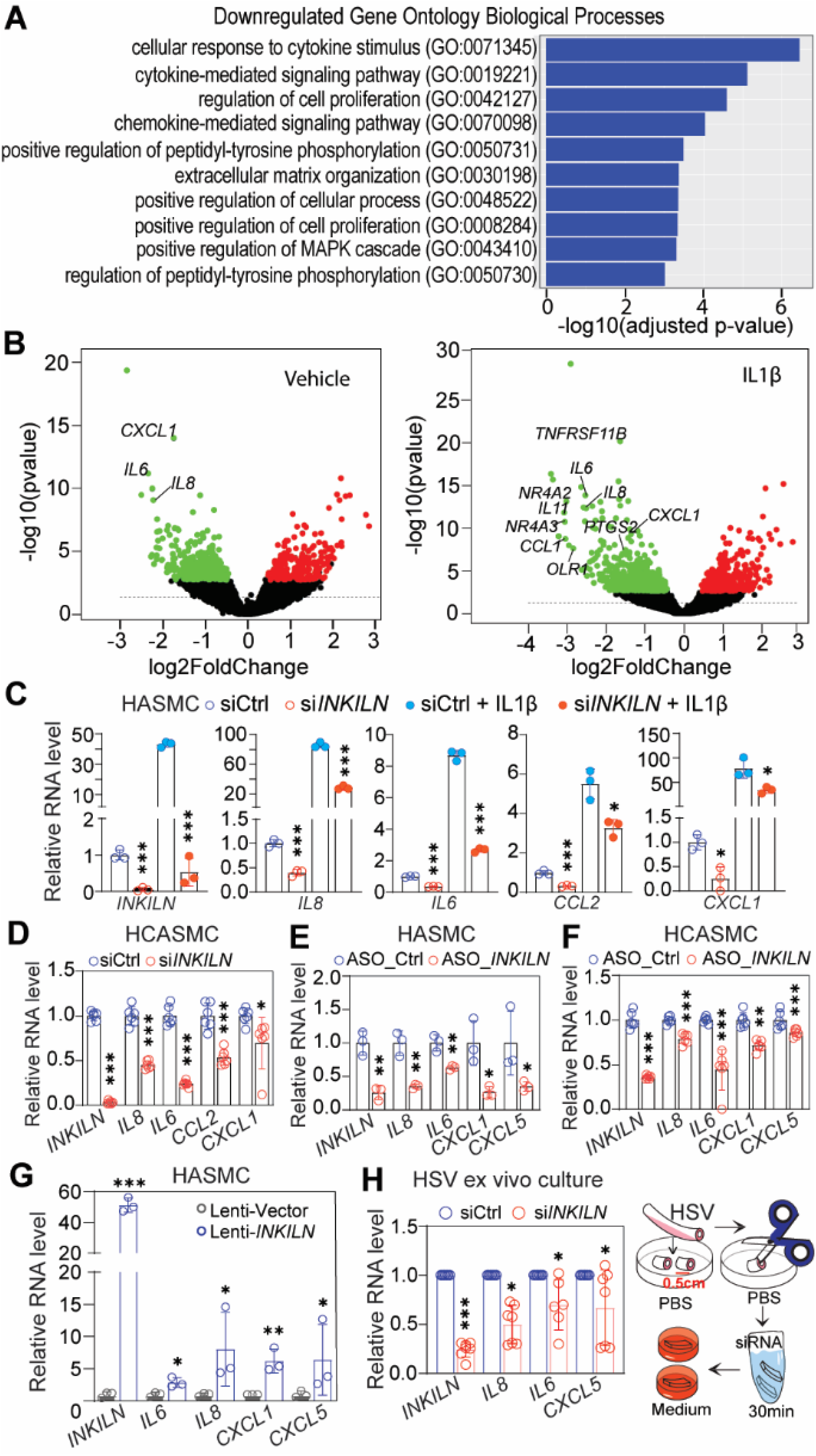
*INKILN* positively regulates proinflammatory gene expression. **A**. The top 10 enriched Gene Ontology (GO) biological process terms downregulated by *siINKILN* in HASMCs under the IL1β-induced condition are shown (adjusted p<0.05 and absolute log2FoldChange). Individual GO terms were sorted by adjusted p values. **B.** Volcano plot depicts the differentially expressed genes in HASMCs ± IL1β treated with si*INKILN* versus siCtrl (sicontrol). **C**-**F.** qRT-PCR validation of the reduced expression of the indicated pro-inflammatory genes upon *INKILN* depletion in HASMCs ± IL1β (**C,** n=3) and growing HCASMCs (**D,** n=6) using *siINKILN* versus siCtrl (n=3) or FANA Antisense Oligonucleotides (ASO) to *INKILN (ASO_INKILN)* versus ASO control (ASO_Ctrl) in growing HASMCs (**E**, n=3) and HCASMCs (**F,** n=3). **G**. Growing HASMCs transduced with the same amount of lentivirus carrying the *INKILN* (Lenti-*INKILN*) or lentivirus negative control (Lenti-vector) for 72 hours before RNA extraction for qRT-PCR of the indicated proinflammatory genes (n=3). **H**. HSV segments incubated with *siINKILN* or siCtrl for 30 minutes at the dose of 25 nM followed by ex vivo culture for 3 days before total RNA isolation for qRT-PCR of the indicated genes (each dot represents the average value from 3 separate segments from the same patient, n=6 patients). Unpaired two-tailed Student’s t test. * p<0.05, ** p<0.01, *** p<0.0001, ns, not significant.

### *INKILN* interacts with MKL1 in the cytoplasm of VSMCs

To gain insight into the mechanism through which *INKILN* promotes inflammatory gene expression, we sought to determine the cellular localization of *INKILN* in cultured VSMCs. qRT-PCR of total RNA from fractionated HCASMs showed that *INKILN* was distributed primarily in the cytosolic compartment (**Figure 4A**), a finding further confirmed by single molecule RNA-FISH (**Figure 4B, left**). *INKILN* signal was authenticated by siRNA-mediated *INKILN* gene knockdown and a negative control probe (**Figure 4B, middle; Supplemental Figure 4A**). Quantitation of *INKILN* positive cells revealed an average copy number of ~17 *INKILN* transcripts per cell (**Figure 4B, right)**. Because of the robust induction of MKL1 protein levels and its established role in vascular inflammation and disease, ^34 36^ and its high RNA binding potential revealed by a well-recognized algorithm, CatRAPID (**Supplemental Figure 4B**), ^57^ we sought to test if MKL1 could be the interactive partner of *INKILN* to activate the proinflammatory gene program. In vitro RNA pulldown coupled with Western blotting revealed enriched interaction between MKL1 and the *INKILN* transcript, but not the negative control corresponding to antisense *INKILN* (**Figure 4C**). This interaction appears specific, as two well-recognized activators of inflammation, p65 and p38, were not pulled down by *INKILN* (**Supplemental Figure 4C**). Further, RNA immunoprecipitation (RIP)-qPCR showed a high enrichment of *INKILN* in the RNA precipitates pulled down by MKL1 antibody, but not p65 antibody and the negative control IgG in HCASMCs (**Figure 4D**) and human rhabdomyosarcoma (RD) cells (**Figure 4E**). This enrichment was specific to *INKILN* as two abundant transcripts, *18S* (**Figure 4D**) and *RNU6-1* (**Figure 4E**) were not enriched by anti-MKL1 precipitation. Immuno-RNA-FISH studies reveal *INKILN* and MKL1 protein mainly colocalize in the cytosol of HCASMs (**Figure 4F**). Such cytosolic colocalization was specific to MKL1 as no colocalization was seen with ACTA2, a highly expressed cytoskeletal protein, and the species-matched negative control IgG in VSMCs **(Figure 4F; Supplemental Figure 4D**). Quantitation of such colocalization through Pearson correlation coefficient analysis revealed a higher correlation score for MKL1 with *INKILN,* but not *PPIB* (**Figure 4G**). To further validate cytosolic colocalization between *INKILN* and MKL1, we stimulated the cells with Jasplakinolide (Jpk) and TGFβ1, two established activators of MKL1 nuclear translocation.^58^ Though enhanced nuclear MKL1 was seen after stimulation by both activators, colocalization was retained in the cytosol (**Figure 4H**, **Left**). The correlation in colocalization between MKL1 and *INKILN* was reduced upon either Jpk or TGFβ1 treatment compared with their individual controls (**Figure 4H, Right**), likely attributable to the decreased amount of cytosolic MKL1 protein following nuclear translocation. These data support *INKILN* physically interacting with MKL1 in the cytosolic compartment of VSMCs.

**Figure 4.**
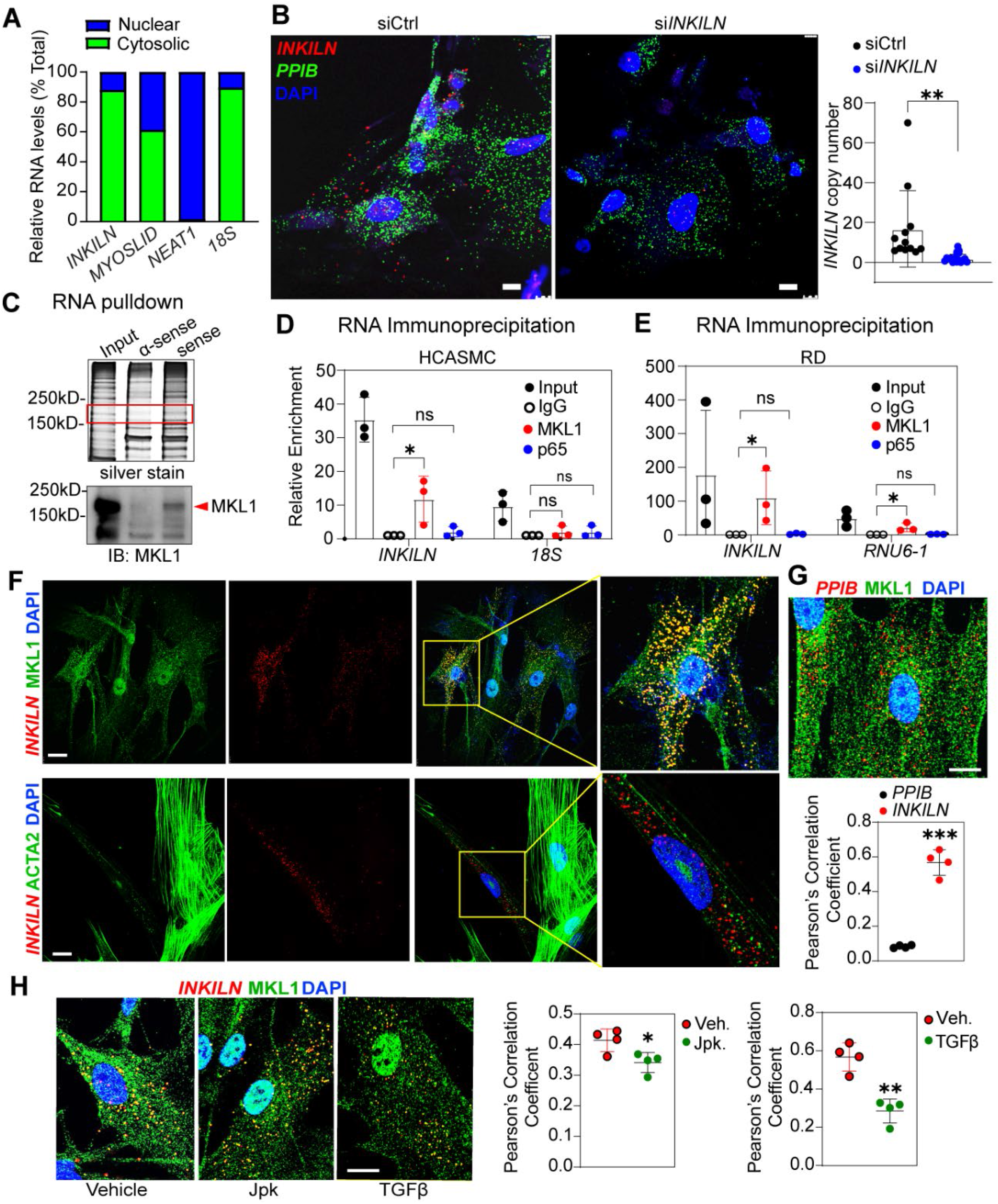
*INKILN* interacts with MKL1 in the cytoplasm of VSMCs. **A**. Representative qRT-PCR analysis of *INKILN* and the indicated control genes in total RNA from the fractionated cytosolic and nuclear compartments in HCASMCs (n=3 independent experiments). **B.** RNA-FISH for *INKILN* (red) and *PPIB* (green) and DAPI (blue) staining in growing HCASMCs and the quantitation of the copy number of *INKILN* per cell (n=12 fields with 39 cells for siCtrl and n=17 fields with 71 cells for *siINKILN* from 3 biological replicates quantitated). **C.** In vitro RNA pulldown using biotinylated sense *INKILN* and antisense *INKILN* RNA showed an enriched band between 150kD and 250kD with sense *INKILN* by silver staining (red rectangle), which was validated as MKL1 protein by western blot (below). Representative images shown (n=3). **D-E.** Representative RNA Immunoprecipitation (RIP)-qPCR in HCASM cells (**D**) and human rhabdomyosarcoma (RD) cells (**E**) showed an enrichment of *INKILN* from RNA precipitates by MKL1, but not p65 antibodies (n=3 independent experiments). **F-G**. Representative immuno-RNA-FISH for *INKILN* (red) and MKL1 protein (green) in HCASMCs (**F**) and the quantitation of the co-localization between *INKILN* and MKL1 protein by Pearson correlation coefficient analysis (**G**). *PPIB* mRNA was used as a negative control which fails to co-localize with MKL1 protein (quantitation was from 4 cells of 1 representative experiment out of 3 independent experiments). **H.** Representative immuno-RNA-FISH for *INKILN* and MKL1 protein in HCASMCs treated with Jasplakinolide (Jpk) for 6 hours or TGFβ for 24 hours to induce MKL1 nuclear translocation, and the quantitation of the co-localization of *INKILN* with MKL1 by Pearson correlation coefficient analysis under both stimulation conditions relative to their individual vehicle controls (quantitation was from 4 cells of 1 representative experiment out of 3 independent experiments). Scale Bar =20μm. **B**, **G**, and **H**, unpaired two-tailed Student’s t test; **D** and **E**, pair matched one-way ANOVA followed by Dunnett’s multiple comparison test. * p<0.05, ** p<0.01, *** p<0.0001, ns, not significant.

### Loss of *INKILN* suppresses MKL1/p65-mediated activation of the proinflammatory gene program

The transcription factor MKL1 functions as a critical activator of inflammation in both cultured VSMCs and vascular disease models.^34, 36^ One well-documented mechanism underlying MKL1 activation of vascular inflammation is through the p65/NF-κB pathway.^34, 36^ Knockdown of *MKL1* via either lentivirus-sh*MKL1* or siRNA pool targeting *MKL1* in HCASMCs attenuated the expression of a battery of proinflammatory genes as well as the phosphorylated form of p65 (p-p65) (**Figure 5A, 5B**). Further, increased p-p65 was seen in HCASMs transduced with Ad-MKL1 **(Figure 5C**). These results prompted us to test if *INKILN* promotes the transactivity of MKL1/p65 on the proinflammatory gene program. To test this hypothesis, we first examined the effect of *INKILN* depletion on IL1β-induced p65 nuclear translocation in HASMCs. Western blot of the fractionated protein lysate revealed that IL1β increased the amount of p65 in the chromatin in HASMCs. Such increase was sharply attenuated by siRNA-mediated *INKILN* knockdown. In contrast, neither IL1β stimulation nor *INKILN* knockdown significantly changed p65 protein levels in cytosol or nucleoplasm of HCASMCs (**Figure 5D**). In line with this, immunostaining showed a reduction of nuclear p65 protein in HCASMCs upon *INKILN* knockdown and IL1β stimulation (**Figure 5E**). Further, siRNA-mediated *INKILN* knockdown prevented IL1β-induced MKL1 nuclear translocation (**Figure 5F**). The immunostaining of MKL1 and p65 was authenticated by the species-matched negative control IgG (**Supplemental Figure 5**). These results suggest that *INKILN* may be critical for the nuclear interaction between MKL1 and p65 to transactivate the proinflammatory gene program. Consistent with this notion, a significant reduction of TNFα-activated NF-κB reporter activity was seen in HASMCs treated with *siINKILN* (**Figure 5G**). Collectively, these results support a positive role for *INKILN* in MKL1/p65 transactivation of the proinflammatory gene program.

**Figure 5.**
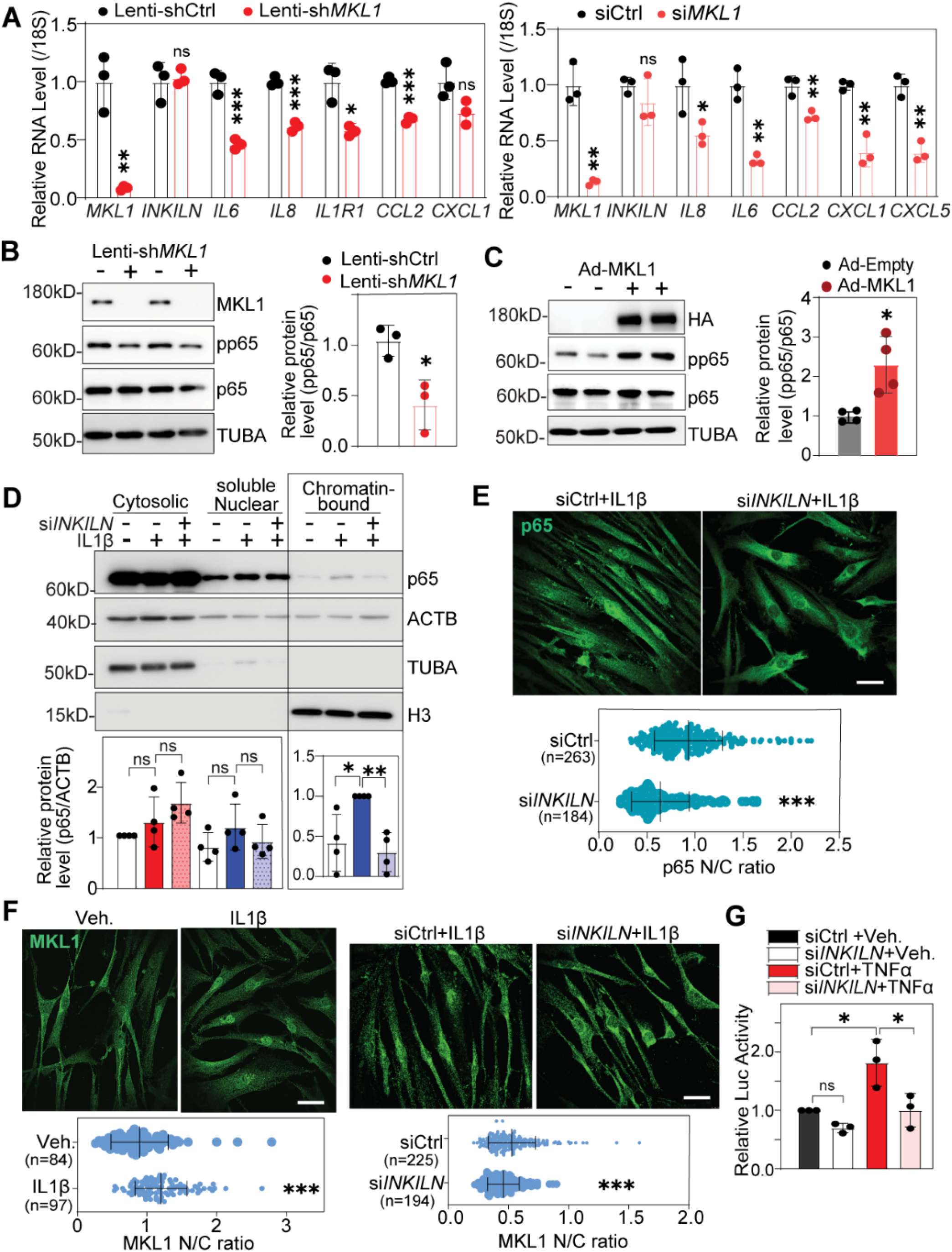
Loss of *INKILN* suppresses MKL1/p65-mediated activation of the proinflammatory gene program. **A.** qRT-PCR analysis of the expression levels of the indicated proinflammatory genes in HCASMCs treated with same amount of lentivirus carrying short hairpin RNA to MKL1 (Lenti-sh*MKL1*) or siRNA SMART POOL to MKL1 *(siMKL1)* versus their individual controls (Lenti-shCtrl or siCtrl) (n=3). **B-C.** Representative western blot of the phosphorylated p65 (pp65) level in HCASMCs transduced with Lenti-sh*MKL1* versus Lenti-shCtrl (**B**, n=3) or Adenovirus carrying MKL1 transcript *(Ad-MKL1)* versus Ad-empty control (Ad-empty) for 48 hours before protein extraction for western blot of the indicated proteins (**C**, n=4) and the respective quantitation. **D**. Representative western blot of fractionated proteins from the indicated cellular compartments in HCASMCs depleted by *siINKILN* for 48 hours followed by IL1β stimulation for 24 hours and the quantitation (n=4). **E.** Representative immunofluorescence staining for p65 protein in HASMCs treated with *siINKILN* or siCtrl for 48 hours prior to IL1β induction for 24 hours and the quantitation (n=3). **F.** Immunofluorescence staining for MKL1 in HASMCs treated with *siINKILN* versus siCtrl for 48 hours followed by IL1β induction for 24 hours (n=3 with indicated total cell numbers). **G.** Luciferase assay for NF-κB reporter activity in HASMCs depleted by *siINKILN* for 48 hours followed by TNFα (10 ng/ml) simulation for 6 hours (n=3). **A, B, C, E, F**, unpaired two-tailed Student’s t test; **D**, one-way ANOVA followed by Dunnett’s multiple comparison test; **G**, two-way ANOVA followed by Sidak multiple comparison test. * p<0.05, ** p<0.01, *** p<0.0001, ns, not significant.

### Loss of *INKILN* reduces MKL1 protein stability via enhancing ubiquitination proteasome degradation

Data above suggest that MKL1/p65 may mediate *INKILN* activation of the proinflammatory gene program in VSMCs. To further delineate the molecular mechanism, we performed co-immunoprecipitation (Co-IP) in HCASMCs to assess if *INKILN* impacts the interaction between p65 and MKL1, an important mechanism underlying the activation of vascular inflammation.^59^ *INKILN* knockdown diminished the association between MKL1 and p65 in HCASMCs. Interestingly, while *INKILN* knockdown displayed no effect on the levels of input p65, it caused a notable reduction of the input MKL1, suggesting *INKILN* may positively regulate MKL1 protein abundance (**Figure 6A**). Indeed, depletion of *INKILN* by two separate siRNAs significantly reduced the protein levels of MKL1 in HCASMCs (**Figure 6B**). This result was reproduced in HASMCs and RD cells (**Figure 6C, 6D**). Knockdown of *INKILN* had no significant effect on *MKL1* mRNA levels (**Figure 6E**), suggesting that the influence of *INKILN* on MKL1 protein abundance was not attributable to changes in *MKL1* transcription or mRNA stability. We next considered whether *INKILN* influences MKL1 protein stability. To test this idea, we first assessed if MKL1 undergoes proteasome degradation in VSMCs. Incubation of HASMC with a proteasome inhibitor, MG132, significantly increased MKL1 protein levels (**Figure 6F**), suggesting that MKL1 was subjected to proteasome-mediated protein degradation in VSMCs as reported previously.^37^ To ascertain if *INKILN* could impact this process, we depleted *INKILN* in HASMCs with siRNA followed by MG132 treatment. MG132 completely rescued the downregulation of MKL1 protein caused by *INKILN* loss in HCASMCs (**Figure 6G**). Finally, immunoprecipitation assays revealed that loss of *INKILN* increased the ubiquitinated form of MKL1 compared with that from control siRNA (**Figure 6H**). These findings suggest that *INKILN* suppresses MKL1 ubiquitination proteasome degradation, leading to the elevated levels of MKL1 protein in VSMCs.

**Figure 6.**
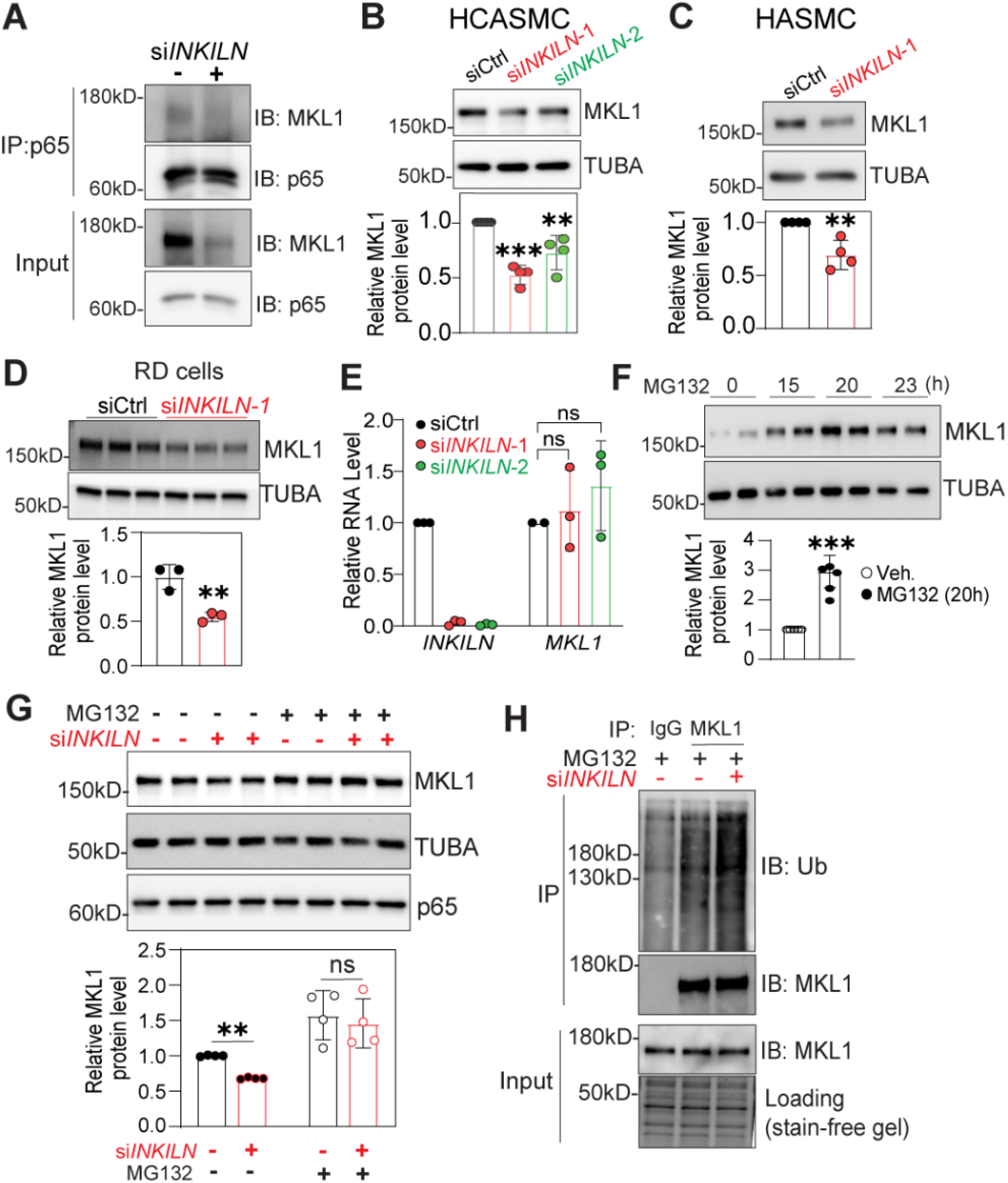
Loss of *INKILN* reduces MKL1 protein stability via enhancing ubiquitination proteasome degradation. **A.** HASMCs were transfected with *siINKILN* or sicontrol for 48 hours prior to protein extraction for p65 immunoprecipitation followed by western blotting analysis of the indicated proteins.1/100 amount of total cell lysates were used as input control. Representative western blot images for the indicated proteins (n=5). **B-D.** HCASMCs (**B**, n=4), HASMCs (**C**, n=4), and RD cells (**D**, n=3) were treated with *siINKILN* or sicontrol for 72 hours and protein lysates were used for western blot analysis of MKL1. Representative western blot images (**B-D**, top) and the quantitation (**B-D**, bottom). **E.** qRT-PCR of *MKL1* expression after siRNA-mediated *INKILN* gene knockdown in HCASMCs (n=3). **F.** HASMCs treated with 5μM MG132 for the indicted time before protein extraction for western blot of MKL1. Representative western blot (top) and the quantitation at 20 hours after the treatment of MG132 (bottom) (n=5). **G**. HASM cells treated with siRNA for 48 hours followed by MG132 (5 μM) for 20 hours prior to protein extraction for western blot of MKL1. Representative western blot image (top) and the quantification (bottom) (n=4). **H.** HCASMCs treated with si*lNKILN* versus sicontrol for 48 hours followed by MG132 (5 μM) treatment for 20 hours prior to protein extraction for immunoprecipitation of MKL1 and western blot of ubiquitin. Representative images were shown (n=4). **B** and **E**, one-way ANOVA followed by Dunnett’s multiple comparison test; **C,D,F**, unpaired two-tailed Student’s t test; **G**, One-way ANOVA followed by Dunnett’s multiple comparison test; unpaired two-tailed Student’s t test. * p<0.05, ** p<0.01, *** p<0.0001, ns, not significant.

### *INKILN* facilitates the interaction between MKL1 and USP10

Ubiquitin proteasome degradation is subjected to the governance of enzyme chains, comprising E1 activating, E2 conjugating, and E3 ligase enzymes, which ultimately leads to the formation of polyubiquitin chains on a target substrate.^60^ This process can be reversed by diverse deubiquitinases (DUBs), notably Ubiquitin Specific Peptidase (USP) family members that cleave ubiquitin from ubiquitin-conjugated protein substrates.^61, 62^ Among all USP family members, USP10 has emerged as a key regulator of critical biological processes, including immune response and inflammation.^63, 64^ To test if USP10 participates in the de-ubiquitination of MKL1, we first examined if MKL1 physically interacts with USP10 in VSMCs. Co-IP showed MKL1 forms a complex with USP10, a finding further supported by colocalization revealed by immunofluorescence staining **(Figure 7A, 7B)**. Further, siRNA-mediated USP10 knockdown decreased, while adenovirus overexpressing USP10 increased, protein levels of MKL1 in HCASMCs **(Figure 7C, 7D)**. These results suggest that USP10 may serve as a critical DUB to inhibit MKL1 ubiquitination proteasome degradation. To determine how *INKILN* participates in the regulation of USP10 on MKL1 protein, we went on to examine the influence of *INKILN* on USP10 gene expression. Analysis of bulk RNA-seq in *INKILN*-depleted HASMCs failed to reveal a consistent effect of *INKILN* knockdown on the levels of *USP10* mRNA (**GSE158219**). Western blot showed a marginal, but statistically significant, reduction of USP10 protein levels upon *INKILN* knockdown (**Figure 7E**). These results suggest that *INKILN* may not have a major role in regulating USP10 expression. We then assessed whether *INKILN* influences the physical interaction between USP10 and MKL1. Co-IP showed that the interaction between MKL1 and USP10 was significantly attenuated by *INKILN* knockdown in HCASMCs **(Figure 7F)**. Proximity ligation assay (PLA) is a well-recognized approach to determine the physical interactions between proteins.^65^ Consistently, PLA revealed a clear interaction between MKL1 and USP10 in HCASMCs, and such interaction was significantly attenuated upon *INKILN* knockdown (**Figure 7G**). These results suggest that *INKILN* acts as a scaffold to facilitate USP10 deubiquitination of MKL1 protein. The immunostaining of USP10 and PLA for USP10/MKL1 were authenticated by *siUSP10* and *siMKL1* (**Supplemental Figure 6A, B**). Similar to MKL1, there is a clear enrichment of *INKILN* from the RNA precipitates against USP10 antibody but not the negative control IgG in HCASMCs. This interaction appeared to be specific, as lncRNA *NEAT1* was not enriched in the RNA precipitates **(Figure 7H)**. Taken together, these results suggest that *INKILN* stabilizes MKL1 protein through scaffolding MKL1 and USP10, leading to suppression of MKL1 ubiquitin proteasome degradation.

**Figure 7.**
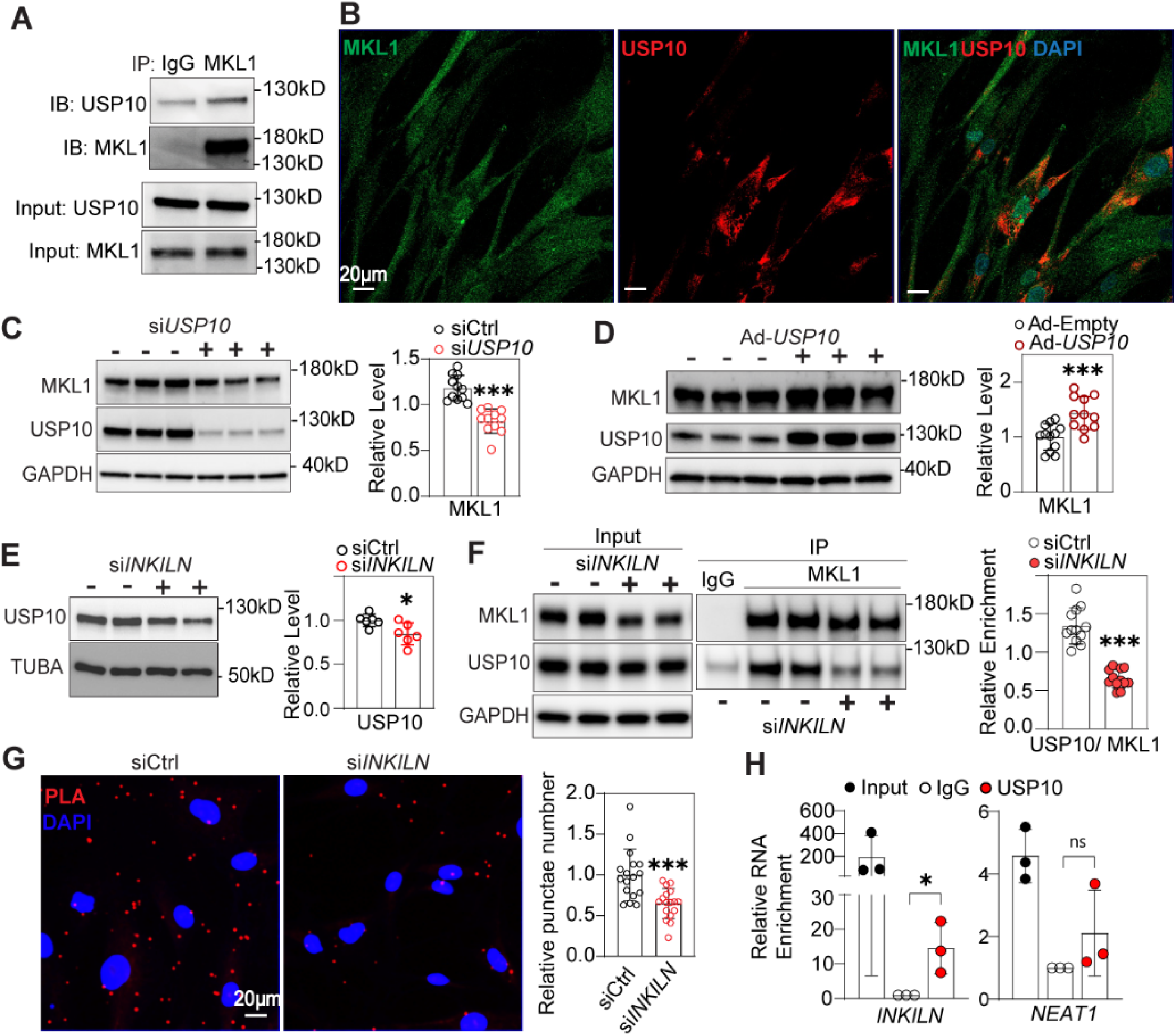
*INKILN* facilitates the interaction between MKL1 and USP10. **A.** Representative image of co-immunoprecipitation of MKL1 followed by western blot of the indicated proteins in HCASMCs (n=4). **B**. Representative co-immunofluorescence staining of MKL1 and USP10 in HCASMCs (n=3). Scale Bar=20μm. **C,D.** Representative western blot of the indicated proteins in HCASMCs treated with siRNA to *USP10* (**C**), or adenovirus overexpressing *USP10* (**D**) versus their individual controls and the quantitation of MKL1 protein levels (n=11). **E.** HCASMCs were treated with siRNA-*INKILN* for 72 hours and the protein levels of USP10 were detected by western blot (n=6). **F.** HCASMCs were treated with siRNA-*INKILN* for 72 hours prior to immunoprecipitation of MKL1 and western blot of the indicated proteins. Representative image was shown and the quantitation was 8 biological replicates from 4 independent experiments. **G**, Representative image of proximity ligation assay (PLA) for MKL1 and USP10 in HCASMCs, and quantitation of PLA punctae was shown (17 fields from n=5 independent experiments). **H.** qRT-PCR of the indicated genes from the RNA pools precipitated by USP10 antibody in HCASMCs (n=3). Unpaired two-tailed Student’s t test. * p<0.05, ** p<0.01, *** p<0.0001, ns, not significant.

### *INKILN* expression is induced in diseased vessels of BAC transgenic mice and promotes ligation injury-induced neointimal formation

Because *INKILN* is a human-specific lncRNA, with no known mouse ortholog, loss-of-function studies in vivo are not possible. To circumvent this limitation, we generated a humanized transgenic mouse strain carrying the *INKILN* and *UMLILO* gene loci and a newly annotated gene *(ENSG00000289530)* found in the first intron of *INKILN* plus upstream and downstream sequences. Importantly, the neighboring protein-coding gene, *IL8*, was deleted through BAC recombineering (**Supplemental Figure 7B; UCSC Genome Browser**).^13^ We genotyped transgenic mice using multiple primers targeting different regions of *INKILN* (**Supplemental Figure 7A**). Using CRISPR-LRS (long read sequencing),^49^ we mapped this transgene as 2 copies on Chr11 of mouse genome at 32,808,215 bp - 32,828,044 bp (mm10), which corroborates qPCR (**Figure 8A, B**).^66^ CRISPR-LRS libraries found mild genome perturbations on the terminal ends of the tandem transgenes comprised of both human and mouse genome sequences (**Supplemental Figure 7B**). To assess proper editing of BAC RP11-997L11, CRISPR-LRS libraries queried inside of the human BAC. Targeting from the 5’ end of *INKILN* through *IL8* found successful and exclusive removal of human *IL8* gene and its proximal promoter. This feature of the *INKILN* BAC mouse allowed us to properly study the function of *INKILN* without the confounding effects of *IL8* expression (**Supplemental Figure 7B**). The only protein-coding gene affected in the mouse genome was *Smim23*, a testis-specific gene of unknown function (**Supplemental Figure 7C**). We refer to this BAC transgenic (Tg) mouse as BAC *INKILN* Tg. qPCR showed that *INKILN* was undetectable in aortas and carotid arteries under physiological conditions, but robustly induced in ex vivo cultured aortas and carotid arteries subjected to complete ligation (**Figure 8C, 8D**). There was no detectable *UMLILO* or ENSG00000289530 gene expression under these conditions (**data not shown**). The injury-induced *INKILN* expression in the carotids was further validated by RNA FISH using a specific probe to *INKILN* (**Figure 8E**). Further, *INKILN* was robustly induced by LPS in BMDMs from *INKILN* Tg mice (**Supplemental Figure 7D**). Finally, immuno-RNA FISH confirmed the colocalization of *INKILN* and mouse MKL1 protein in LPS-induced BMDMs isolated from *INKILN* Tg mice, suggesting a similar physical interaction between *INKILN* and MKL1 in BAC *INKILN* Tg mice (**Supplemental Figure 7E**). These results are congruent with the induction of *INKILN* in phenotypically modulated human SMCs, and in atherosclerotic and aneurysmal human vessels.

**Figure 8.**
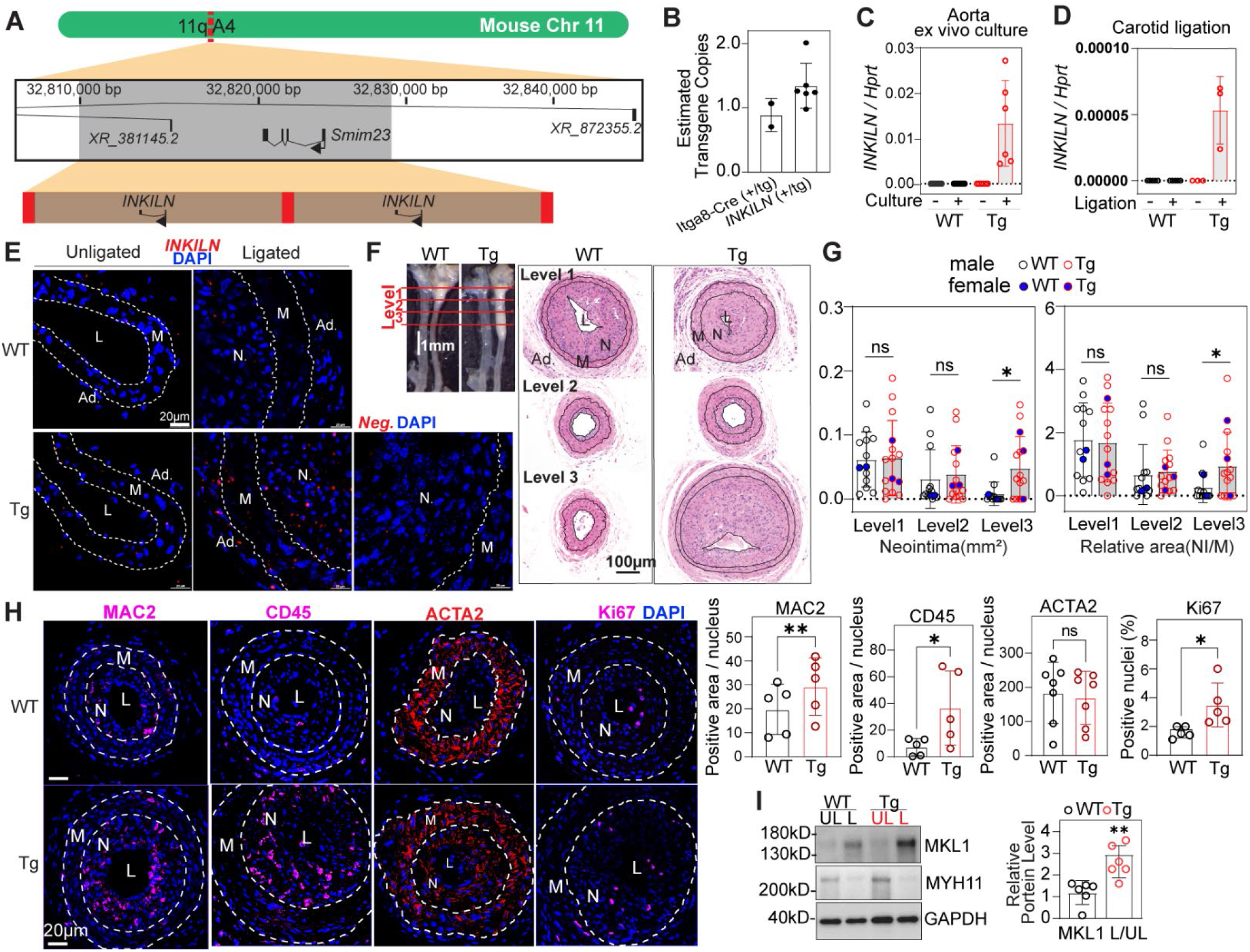
*INKILN* expression in *INKILN* BAC transgenic mice and its influence on neointimal formation. **A.** CRISPR-LRS mapped a single integration locus for human *INKILN.* The integration locus, indicated by a grey box, spanned 32,808,215bp - 32,828,044bp on mouse chromosome 11 (mm10), disrupting testis-specific protein-coding gene *Smim23. INKILN* BAC transgenes (brown rectangles), integrated in a tandem head to tail fashion accompanied with BAC cloning vector sequence (red boxes). **B.** qPCR determined ~2 transgene copies for human *INKILN* (+/tg, n=6) with data normalized to internal control locus. *Itga8-CreER^T2^* mice ^2^ (n=2) served as calibrator for one copy of a transgene. Values graphed as mean ± SEM. **C.** qRT-PCR of *INKLIN* for the uncultured versus 3 days ex vivo cultured aorta segments from WT and *INKILN* transgenic (Tg) mice (n=6). **D.** qRT-PCR of *INKLIN* for unligated versus 1 week ligated carotid arteries from WT and Tg mice (n=3). **E**. Representative RNA FISH image for *INKILN* transcripts in unligated versus 4 week ligated carotid arteries from WT and Tg mice (n=3). **F, G.** Representative whole mount of 4 week ligated carotid arteries from WT versus Tg mice (**F**, left), the H&E staining of sections at different levels (**F**, right), and the quantitation of neointimal formation (**G**, n=13 for WT and n=15 for Tg). Representative images of immunofluorescence staining (**H**) for the indicated proteins on cross sections of ligated carotid arteries from WT and Tg mice and the quantitation of the fluorescence positive area over the total nuclei at neointima and media (n= 5 mice, 1 section/mouse at level 3). **I**. Western blot of the indicated proteins in unligated and ligated carotid arteries from WT versus *INKILN* Tg mice and the quantitation (n=6). NI, Neointima; M, Media; UL, unligated carotids; L, ligated carotids. **G**, **H**, **I**, paired two-tailed Student’s t test. *p<0.05, **p<0.01, ns, not significant.

Next, we sought to determine the influence of *INKILN* on ligation injury-induced neointimal formation in *INKILN* Tg mice. We subjected *INKILN* Tg and littermate WT control mice to left carotid artery ligation injury for 3 weeks. H&E staining of the serial sections at defined distances from the ligation suture site showed significantly increased neointimal formation in the distal region (level 3) of *INKILN* Tg mice relative to littermate WT control mice; comparable neointimal formation was seen in level 1 and level 2 regions of *INKILN* Tg and littermate WT control mice (**Figure 8F, 8G**). No discernable difference was observed in the medial layer of injured and sham control carotids in *INKILN* Tg versus WT mice (**Supplemental Figure 7F**). Immunostaining showed increased proinflammatory cell infiltration and cell proliferation, with increased cell numbers positive for macrophage marker MAC2, leukocytes marker CD45, and proliferation marker KI67, respectively, in the injured carotids of *INKILN* Tg mice relative to WT controls (**Figure 8H**). We did not observe significant changes in ACTA2 positive cells in the injured vessels from Tg versus WT mice (**Figure 8H**). Authentication of MAC2 and CD45 staining was conducted using species-matched IgG controls (**Supplemental Figure 7G**). Finally, ligation injured carotids exhibited significantly higher levels of MKL1 protein compared with those in WT mice, suggesting a similar effect of *INKILN* on MKL1 protein stability in *INKILN* Tg mice (**Figure 8I**). These results demonstrate that *INKILN* is induced in the context of vascular injury and exacerbates neointimal formation, consistent with results seen in human cells and vessels.

## Discussion

The present study provides insight into the transcriptional control, proinflammatory role, and mechanistic action of a novel, human-specific lncRNA, *INKILN,* in VSMCs. Evidence is provided for *INKILN* downregulation in differentiated contractile VSMCs and upregulation upon proinflammatory stimuli in a p65/NFkB-dependent manner. *INKILN* activates the proinflammatory gene program in multiple primary VSMCs and ex vivo HSV cultures, and promotes injury-induced neointimal formation in BAC *INKILN* transgenic mice. The molecular basis underlying *INKILN* activation of VSMC inflammation involves its action as a scaffold with MKL1, a major transcriptional activator of vascular inflammation, and the deubiquitinase USP10, which inhibits MKL1 ubiquitination and proteasomal degradation (**Figure 9**). Our study not only uncovers a novel lncRNA activating the VSMC proinflammatory phenotype, but also elucidates a previously unknown pathway governing MKL1 protein stability to potentiate its proinflammatory role. Given the increased recognition of VSMC phenotypic switching to macrophage-like proinflammatory phenotype in the initiation and aggravation of vascular diseases, our study provides important insights into potential therapeutic strategies for vascular diseases via effectively targeting the interplay between coding and noncoding pathways. Because the induction and function of *INKILN* in BAC Tg mice recapitulate those in human cells and tissues, our study also indicates that human BAC Tg mouse models may offer an innovative approach to studying human-specific lncRNAs in the vascular system.

**Figure 9.**
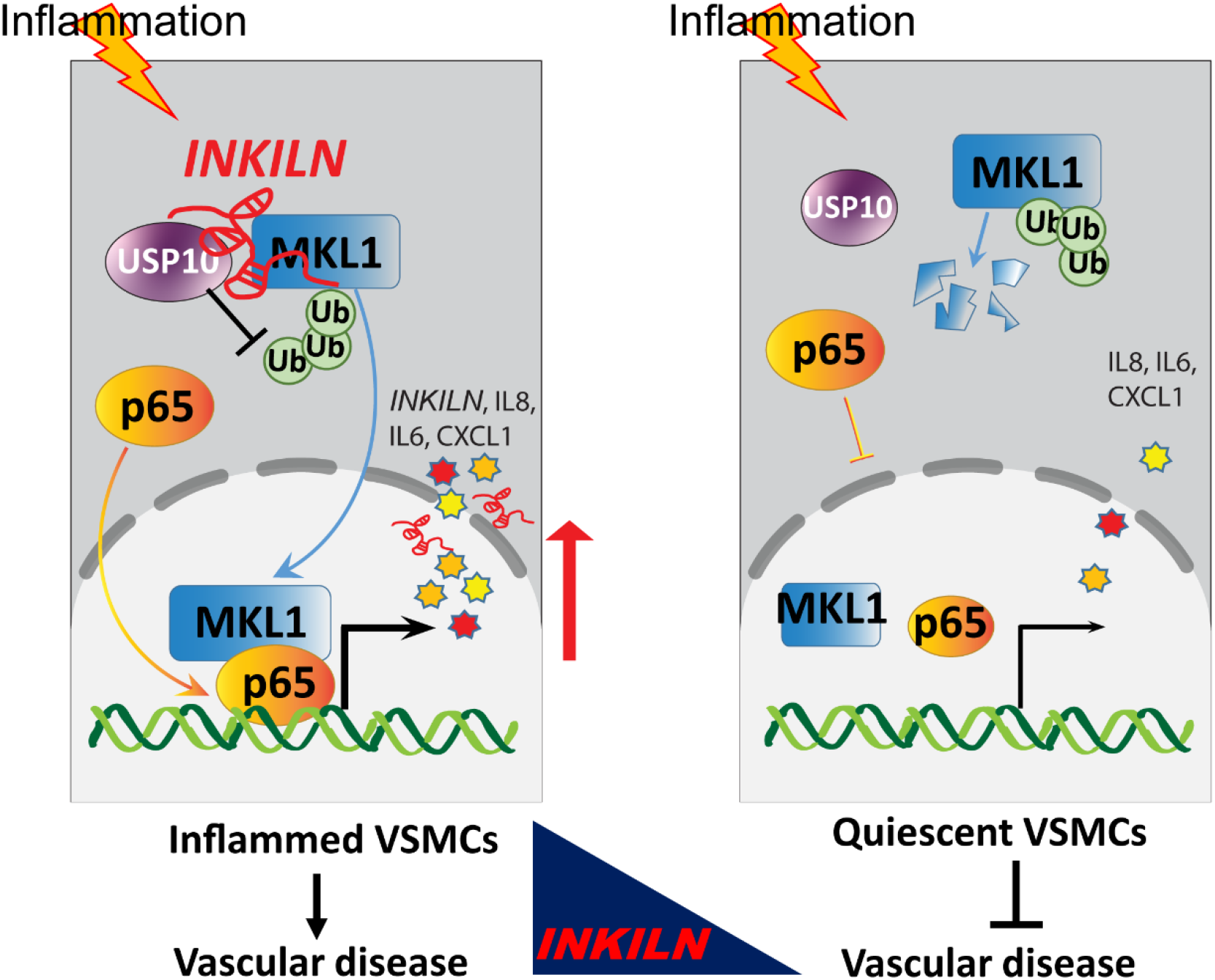
Working model of *INKILN* activating VSMC inflammation. Inflammation induces *INKILN* expression, which inhibits MKL1 ubiquitin proteasome degradation via USP10 and enhances both MKL1 and p65 nuclear translocation, resulting in the increased nuclear interaction of MKL1 with p65 and subsequent transactivation of the proinflammatory gene program.

The genomic localization of *INKILN* is particularly unique, with its first intron harboring the upstream master lncRNA of the inflammatory chemokine locus *(UMLILO)* transcribed in a reverse direction. *UMLILO* is an enhancer lncRNA that facilitates H3K4me3 epigenetic priming of chemokine genes and trained immunity.^67^ UMLILO is barely detectable in our VSMC systems (data not shown). This suggests that the functional role of *INKILN* may be independent of *UMLILO.* It should be noted, though *INKILN* and *UMLILO* reside in the same enhancer RNA region on chromosome 4 and both activate proinflammatory genes, several distinctions exist. First, no overlapping sequence is seen in their annotated transcripts. Second, *INKILN* loss-of-function has no effect on *UMLILO* expression (**Supplemental Figure 8**). Third, *INKILN* is predominately located in the cytosolic compartment whereas *UMLILO* resides in the nucleus. Finally, *UMLILO* activates chemokine genes in *cis*, whose activity is confined to a *CXCL* topologically associated domain.^67^ In contrast, *INKILN* functions in the cytosol and influences a broad range of proinflammatory genes. Therefore, though regulatory roles exerted by these two human-specific lncRNAs appear similar in triggering inflammation, they utilize distinct pathways to achieve functional consequences. Among all lncRNAs annotated in the human genome, the majority of them are restricted to humans.^11^ This is particularly true for lncRNAs identified as modulators of immune and inflammatory responses, including previously published *NKILA, lncRNA-CCL2, INCR1,* and *LUCAT1*.^23, 68–70^ These human-specific immune and inflammatory regulatory lncRNAs could underlie the complexity of immune responses and inflammatory diseases occurring in humans, but not in rodents.

The nearest neighboring protein coding gene to the *INKILN* locus is *IL8,* which is located 20 kb upstream of *INKILN* with no intervening annotated genes. We thus consider *INKILN* and *IL8* as a coding and noncoding gene pair. As a human specific prototypic chemokine, IL8 plays a vital role in inflammation initiation and immune cell chemotaxis, contributing to the pathogenesis of various inflammatory diseases, such as infectious disease, chronic obstructive pulmonary disease, asthma, and cancer.^71–74^ Therefore, inhibition of *IL8* gene expression has emerged as an appealing strategy for therapies against these diseases.^72, 75, 76^ Previous studies reported that *IL8* gene activation involves three distinct regulatory mechanisms: (1) de-repression of the promoter operated by multiple repressors, such as NF-κB-repressing factor, octamer-1 (OCT-1), and HDAC1; (2) transactivation by NF-κB and AP1; and (3) mRNA stabilization by the p38MAPK pathway.^77^ In our current study, we showed that *INKILN* positively regulates *IL8* gene expression, which is consistent with our recent survey wherein a connection between *IL8* and *INKILN* was suggested.^78^ This finding together with the epigenetic activation of *IL8* by the enhancer lncRNA *UMLILO,* implies a new lncRNA-mediated regulatory mechanism underlying *IL8* gene activation, which will have important implications for effectively targeting *IL8* gene expression for therapy. Elegant studies from Dr. Wang’s lab, using an innovative transgenic mouse model, which carries a 166 kilobase BAC encompassing the entire human *IL8* gene locus, have reported a crucial role for IL8 in aggravating inflammation and gastrointestinal tumor formation.^79^ Of note, beyond *IL8*, this BAC also harbors the gene loci of *INKILN, UMLILO,* and *ENSG00000289530.* Given the genomic complexity of the BAC insert, the influence of each of these gene products on phenotypes seen in BAC transgenic mice should be taken into account. Finally, careful analysis of the chromatin landscape of *INKILN* and *IL8* revealed multiple peaks of active epigenetic markers, such as H3K27ac and H3K4Me1, suggesting potential enhancers that participate in the transactivation of both genes **(Supplementary Figure 2A)**. The transcription of both *INKILN* and *IL8* is likely subjected to their individual NF-κB site(s) identified in their proximal promoter regions reported here and in previous studies.^77^ Elucidating the in vivo functional role of these regulatory sites awaits future investigative work using genome editing tools as shown for other control elements effecting lncRNA gene expression.^46, 47^

It has long been recognized that the action of MKL1 on transcriptional regulation is signal-responsive and tightly modulated by actin dynamics involving its nucleocytoplasmic translocation.^80^ In addition, recent studies have reported that MKL1 can also modulate gene expression through epigenetic pathways, mainly by interacting with multiple histone modifiers, including Brahma-related gene-1 (BRG1), COMPASS/COMPASS-like complex, and WD Repeat Domain 5 (WDR5), to ensure an active chromatin status for gene transcription.^35, 81, 82^ Compared with these well-established mechanisms underlying MKL1 transactivity, the regulation of MKL1 expression, especially at the protein level, is limited.^37^ Here, we report on a novel paradigm for MKL1 protein stability, which involves the coordinated actions of *INKILN* and the deubiquitinase USP10 in the cytoplasm. The fact that *INKILN* depletion results in a decreased interaction between MKL1 and USP10 suggests that *INKILN* acts as a scaffold for USP10 and MKL1 to facilitate USP10-mediated deubiquitination of MKL1. Given the increased levels of *INKILN* expression reported here and MKL1 protein under vascular disease contexts such as aortic dissection and aneurysm,^33, 34^ we propose *INKILN* serves as a previously unknown mechanism underlying MKL1 protein stability during disease progression. The detailed mechanism as to how the regulatory axis of *INKILN*/USP10/MKL1 operates to stabilize MKL1 protein awaits further investigation.

One distinct paradigm derived from our current study is utilization of human BAC Tg mice for elucidating the vascular disease-associated regulation and function of human-specific non-conserved lncRNAs. The majority of transcribed lncRNAs in the human genome are human-specific, lacking orthologs in rodents. Bioinformatics studies suggest that up to 2/3 of non-conserved human-specific long intergenic ncRNAs (lincRNAs) are associated with cardiometabolic traits.^12^ In vivo characterization of those human-specific lncRNAs in a physiological manner represents a big challenge. In our current study, we generated a human BAC which carries the intact *INKILN* gene locus and surrounding sequences, providing a physiologically relevant genomic milieu for *INKILN* gene transcription. The gene expression pattern of *INKILN* in Tg mice mirrors *INKILN* in human cells and tissues, indicating the human BAC Tg mouse captures critical regulatory elements required for *INKILN* transcription. Notably, we observed a significant increase in neointimal formation in *INKILN* Tg mice, which is in line with the proinflammatory function of *INKILN* in humans. These results suggest that humanized BAC Tg mice may offer a physiologically relevant tool for in vivo investigation of human specific lncRNAs, whose studies currently are largely confined to in vitro cultured cells. To our knowledge, this is the first report harnessing human BAC Tg mice to investigate the function of a non-conserved human lncRNA in the vascular system.

Several lines of key evidence are provided to support the proinflammatory role of *INKILN* in VSMCs. First, *INKILN* is highly induced in cultured VSMCs by different proinflammatory stimuli, ex vivo cultured HSV segments, and human aneurysm samples. Second, both loss-of-function and gain-of-function studies consistently revealed a positive role for *INKILN* in activating a repertoire of proinflammatory genes. Third, loss of *INKILN* reduced the interaction between p65 and MKL1 protein and the transactivity of p65/NF-κB. These data collectively suggest a novel regulatory axis comprising *INKILN,* MKL1, and p65 to potentiate the proinflammatory gene program in VSMCs. The attenuation of MKL1 nuclear translocation upon *INKILN* depletion is intriguing, and may be partially attributive to the decreased MKL1 protein pool. In addition, beyond the cytosolic interaction with USP10 and MKL1 to stabilize MKL1 protein, *INKILN* could sequester away G-actin for MKL1 binding, thereby releasing MKL1 for nuclear shuttling. On the other hand, though the reduced p65 nuclear translocation was consistently seen in VSMCs with *INKILN* knockdown, we could not detect the physical association between *INKILN* and p65 in our system. We thus surmise that the impact of *INKILN* on p65 nuclear translocation is likely indirect. One possibility is the influence of the proinflammatory cytokines and chemokines that are activated by *INKILN.*

There are several limitations in our current study. First, the NF-κB site we defined only exhibited moderate response to TNFα treatment. Given the robust activation of *INKILN* gene expression in response to TNFα, additional *cis* element(s) may be responsible for *INKILN* gene transcription. Second, additional mechanisms may underlie *INKILN*-induced activation of proinflammatory gene expression. Third, despite repeated attempts, the physical interaction between *INKILN* and MKL1/USP10 in patient samples and animal models is lacking. Newer assays and reagents will be necessary to overcome the technical challenges in demonstrating such in vivo complexes. Last, there are two limitations with our *INKILN* Tg studies. Though expression of *UMLILO* and a newly annotated gene *ENSG00000289530* are undetectable in vascular cells and Tg mice under conditions of *INKILN* expression (data not shown), the influence on the phenotype of Tg mice through an indirect pathway, for example via cis regulatory pathway, cannot be excluded. Finally, precise elucidation of *INKILN* gene transcription using genome editing approaches cannot be readily conducted with current *INKILN* Tg mice because of two tandem copies of *INKILN*.

In summary, we report the novel lncRNA *INKILN* as a potent activator of VSMC inflammation. The proinflammatory action of *INKILN* is, at least partially, mediated by scaffolding MKL1 and USP10, thereby alleviating MKL1 ubiquitination-mediated proteasome degradation. *INKILN* is the first lncRNA identified that interacts with and stabilizes MKL1 protein to activate VSMC inflammation. Given the emerging roles of VSMC inflammation, and well-recognized role of the VSMC phenotypic switching to macrophage-like transition in the etiology of different vascular disorders, our findings provide new insights into a therapeutic strategy for vascular disease via effectively targeting the interplay between coding and noncoding pathways. Our studies also indicate that human BAC Tg mouse models offer a novel approach for in vivo investigation of human specific lncRNAs in a physiological relevant genomic context.

## Acknowledgements

We thank Rochester Genomics Research Center for performing the RNA-seq experiments and Dr. Deyou Zheng from Albert Einstein College of Medicine for bioinformatics analysis. We thank Mrs. Diane Singer at Albany Medical College for providing us with the primary VSMC cultures.

## Author contributions

WZ, JZ, DL, MMB, and XL designed and performed the research. WZ and XL wrote the paper. WZ, JZ, DL, WW, SS, NI, JP, WK, AWT, YWL, MDB, MB, JR, DK, QL, GW, PG, and MC performed experiments and analyzed the data. HWK, NW, CM, AHB, and LM contributed to human sample studies. JMM, AHB, and LM participated in research design and edited the manuscript. The authors declare no conflicts of interest.

## References

1. Zanoli, L, Briet, M, Empana, JP, Cunha, PG, Mäki-Petäjä, KM, Protogerou, AD, et al. (2020). Vascular consequences of inflammation: a position statement from the ESH Working Group on Vascular Structure and Function and the ARTERY Society. J Hypertens 38: 1682–1698.

2. Shah, PK (2003). Inflammation, neointimal hyperplasia, and restenosis: as the leukocytes roll, the arteries thicken. Circulation 107: 2175–2177.

3. Williams, JW, Huang, LH, and Randolph, GJ (2019). Cytokine Circuits in Cardiovascular Disease. Immunity 50: 941–954.

4. Liberale, L, Ministrini, S, Carbone, F, Camici, GG, and Montecucco, F (2021). Cytokines as therapeutic targets for cardio-and cerebrovascular diseases. Basic Res Cardiol 116: 23.

5. Nguyen, MT, Fernando, S, Schwarz, N, Tan, JT, Bursill, CA, and Psaltis, PJ (2019). Inflammation as a Therapeutic Target in Atherosclerosis. J Clin Med 8.

6. Libby, P, and Everett, BM (2019). Novel Antiatherosclerotic Therapies. Arteriosclerosis, thrombosis, and vascular biology 39: 538–545.

7. Baylis, RA, Gomez, D, Mallat, Z, Pasterkamp, G, and Owens, GK (2017). The CANTOS Trial: One Important Step for Clinical Cardiology but a Giant Leap for Vascular Biology. Arteriosclerosis, thrombosis, and vascular biology 37: e174–e177.

8. Kosmas, CE, Silverio, D, Sourlas, A, Montan, PD, Guzman, E, and Garcia, MJ (2019). Anti-inflammatory therapy for cardiovascular disease. Ann Transl Med 7: 147.

9. Ridker, PM (2019). Anti-inflammatory therapy for atherosclerosis: interpreting divergent results from the CANTOS and CIRT clinical trials. J Intern Med 285: 503–509.

10. Guo, H, Ahmed, M, Zhang, F, Yao, CQ, Li, S, Liang, Y, et al. (2016). Modulation of long noncoding RNAs by risk SNPs underlying genetic predispositions to prostate cancer. Nat Genet 48: 1142–1150.

11. Ruan, X, Li, P, Chen, Y, Shi, Y, Pirooznia, M, Seifuddin, F, et al. (2020). In vivo functional analysis of non-conserved human lncRNAs associated with cardiometabolic traits. Nat Commun 11: 45.

12. Foulkes, AS, Selvaggi, C, Cao, T, O’Reilly, ME, Cynn, E, Ma, P, et al. (2021). Nonconserved Long Intergenic Noncoding RNAs Associate With Complex Cardiometabolic Disease Traits. Arteriosclerosis, thrombosis, and vascular biology 41: 501–511.

13. Long, X, Slivano, OJ, Cowan, SL, Georger, MA, Lee, TH, and Miano, JM (2011). Smooth muscle calponin: an unconventional CArG-dependent gene that antagonizes neointimal formation. Arteriosclerosis, thrombosis, and vascular biology 31: 2172–2180.

14. Andersen, RE, Hong, SJ, Lim, JJ, Cui, M, Harpur, BA, Hwang, E, et al. (2019). The Long Noncoding RNA Pnky Is a Trans-acting Regulator of Cortical Development In Vivo. Developmental cell 49: 632–642.e637.

15. Statello, L, Guo, CJ, Chen, LL, and Huarte, M (2021). Gene regulation by long non-coding RNAs and its biological functions. Nature reviews Molecular cell biology 22: 96–118.

16. Jaé, N, and Dimmeler, S (2020). Noncoding RNAs in Vascular Diseases. Circulation research 126: 1127–1145.

17. Zhao, J, Zhang, W, Lin, M, Wu, W, Jiang, P, Tou, E, et al. (2016). MYOSLID Is a Novel Serum Response Factor-Dependent Long Noncoding RNA That Amplifies the Vascular Smooth Muscle Differentiation Program. Arteriosclerosis, thrombosis, and vascular biology 36: 2088–2099.

18. Leisegang, MS, Fork, C, Josipovic, I, Richter, FM, Preussner, J, Hu, J, et al. (2017). Long Noncoding RNA MANTIS Facilitates Endothelial Angiogenic Function. Circulation 136: 65–79.

19. Das, S, Zhang, E, Senapati, P, Amaram, V, Reddy, MA, Stapleton, K, et al. (2018). A Novel Angiotensin II-Induced Long Noncoding RNA Giver Regulates Oxidative Stress, Inflammation, and Proliferation in Vascular Smooth Muscle Cells. Circulation research 123: 1298–1312.

20. Haemmig, S, Yang, D, Sun, X, Das, D, Ghaffari, S, Molinaro, R, et al. (2020). Long noncoding RNA SNHG12 integrates a DNA-PK-mediated DNA damage response and vascular senescence. Science translational medicine 12.

21. Monteiro, JP, Rodor, J, Caudrillier, A, Scanlon, JP, Spiroski, AM, Dudnakova, T, et al. (2021). MIR503HG Loss Promotes Endothelial-to-Mesenchymal Transition in Vascular Disease. Circulation research 128: 1173–1190.

22. Lyu, Q, Xu, S, Lyu, Y, Choi, M, Christie, CK, Slivano, OJ, et al. (2019). SENCR stabilizes vascular endothelial cell adherens junctions through interaction with CKAP4. Proceedings of the National Academy of Sciences of the United States of America 116: 546–555.

23. Khyzha, N, Khor, M, DiStefano, PV, Wang, L, Matic, L, Hedin, U, et al. (2019). Regulation of CCL2 expression in human vascular endothelial cells by a neighboring divergently transcribed long noncoding RNA. Proceedings of the National Academy of Sciences of the United States of America 116: 16410–16419.

24. Simion, V, Zhou, H, Pierce, JB, Yang, D, Haemmig, S, Tesmenitsky, Y, et al. (2020). LncRNA VINAS regulates atherosclerosis by modulating NF-κB and MAPK signaling. JCI insight 5.

25. Cremer, S, Michalik, KM, Fischer, A, Pfisterer, L, Jaé, N, Winter, C, et al. (2019). Hematopoietic Deficiency of the Long Noncoding RNA MALAT1 Promotes Atherosclerosis and Plaque Inflammation. Circulation 139: 1320–1334.

26. Chen, Y, Zhao, X, and Wu, H (2021). Transcriptional Programming in Arteriosclerotic Disease: A Multifaceted Function of the Runx2 (Runt-Related Transcription Factor 2). Arteriosclerosis, thrombosis, and vascular biology 41: 20–34.

27. Wang, DZ, Li, S, Hockemeyer, D, Sutherland, L, Wang, Z, Schratt, G, et al. (2002). Potentiation of serum response factor activity by a family of myocardin-related transcription factors. Proceedings of the National Academy of Sciences of the United States of America 99: 14855–14860.

28. Du, KL, Chen, M, Li, J, Lepore, JJ, Mericko, P, and Parmacek, MS (2004). Megakaryoblastic leukemia factor-1 transduces cytoskeletal signals and induces smooth muscle cell differentiation from undifferentiated embryonic stem cells. J Biol Chem 279: 17578–17586.

29. Parmacek, MS (2007). Myocardin-related transcription factors: critical coactivators regulating cardiovascular development and adaptation. Circulation research 100: 633–644.

30. Minami, T, Kuwahara, K, Nakagawa, Y, Takaoka, M, Kinoshita, H, Nakao, K, et al. (2012). Reciprocal expression of MRTF-A and myocardin is crucial for pathological vascular remodelling in mice. Embo j 31: 4428–4440.

31. An, J, Naruse, TK, Hinohara, K, Soejima, Y, Sawabe, M, Nakagawa, Y, et al. (2019). MRTF-A regulates proliferation and survival properties of pro-atherogenic macrophages. Journal of molecular and cellular cardiology 133: 26–35.

32. Chen, D, Yang, Y, Cheng, X, Fang, F, Xu, G, Yuan, Z, et al. (2015). Megakaryocytic leukemia 1 directs a histone H3 lysine 4 methyltransferase complex to regulate hypoxic pulmonary hypertension. Hypertension 65: 821–833.

33. Ito, S, Hashimoto, Y, Majima, R, Nakao, E, Aoki, H, Nishihara, M, et al. (2020). MRTF-A promotes angiotensin II-induced inflammatory response and aortic dissection in mice. PloS one 15: e0229888.

34. Gao, P, Gao, P, Zhao, J, Shan, S, Luo, W, Slivano, OJ, et al. (2021). MKL1 cooperates with p38MAPK to promote vascular senescence, inflammation, and abdominal aortic aneurysm. Redox biology 41: 101903.

35. Yang, Y, Chen, D, Yuan, Z, Fang, F, Cheng, X, Xia, J, et al. (2013). Megakaryocytic leukemia 1 (MKL1) ties the epigenetic machinery to hypoxia-induced transactivation of endothelin-1. Nucleic Acids Res 41: 6005–6017.

36. Yu, L, Fang, F, Dai, X, Xu, H, Qi, X, Fang, M, et al. (2017). MKL1 defines the H3K4Me3 landscape for NF-κB dependent inflammatory response. Sci Rep 7: 191.

37. Hinson, JS, Medlin, MD, Taylor, JM, and Mack, CP (2008). Regulation of myocardin factor protein stability by the LIM-only protein FHL2. American journal of physiology Heart and circulatory physiology 295: H1067–h1075.

38. Wu, W, Zhang, W, Choi, M, Zhao, J, Gao, P, Xue, M, et al. (2019). Vascular smooth muscle-MAPK14 is required for neointimal hyperplasia by suppressing VSMC differentiation and inducing proliferation and inflammation. Redox biology 22: 101137.

39. Mahmoud, AD, Ballantyne, MD, Miscianinov, V, Pinel, K, Hung, J, Scanlon, JP, et al. (2019). The Human-Specific and Smooth Muscle Cell-Enriched LncRNA SMILR Promotes Proliferation by Regulating Mitotic CENPF mRNA and Drives Cell-Cycle Progression Which Can Be Targeted to Limit Vascular Remodeling. Circulation research 125: 535–551.

40. Turner, AW, Hu, SS, Mosquera, JV, Ma, WF, Hodonsky, CJ, Wong, D, et al. (2022). Single-nucleus chromatin accessibility profiling highlights regulatory mechanisms of coronary artery disease risk. Nat Genet 54: 804–816.

41. Corces, MR, Trevino, AE, Hamilton, EG, Greenside, PG, Sinnott-Armstrong, NA, Vesuna, S, et al. (2017). An improved ATAC-seq protocol reduces background and enables interrogation of frozen tissues. Nature methods 14: 959–962.

42. Granja, JM, Corces, MR, Pierce, SE, Bagdatli, ST, Choudhry, H, Chang, HY, et al. (2021). ArchR is a scalable software package for integrative single-cell chromatin accessibility analysis. Nat Genet 53: 403–411.

43. Wirka, RC, Wagh, D, Paik, DT, Pjanic, M, Nguyen, T, Miller, CL, et al. (2019). Atheroprotective roles of smooth muscle cell phenotypic modulation and the TCF21 disease gene as revealed by single-cell analysis. Nature medicine 25: 1280–1289.

44. Long, X, and Miano, JM (2011). Transforming growth factor-beta1 (TGF-beta1) utilizes distinct pathways for the transcriptional activation of microRNA 143/145 in human coronary artery smooth muscle cells. J Biol Chem 286: 30119–30129.

45. Fasolo, F, Jin, H, Winski, G, Chernogubova, E, Pauli, J, Winter, H, et al. (2021). Long Noncoding RNA MIAT Controls Advanced Atherosclerotic Lesion Formation and Plaque Destabilization. Circulation 144: 1567–1583.

46. Gao, P, Lyu, Q, Ghanam, AR, Lazzarotto, CR, Newby, GA, Zhang, W, et al. (2021). Prime editing in mice reveals the essentiality of a single base in driving tissue-specific gene expression. Genome biology 22: 83.

47. Choi, M, Lu, YW, Zhao, J, Wu, M, Zhang, W, and Long, X (2020). Transcriptional control of a novel long noncoding RNA Mymsl in smooth muscle cells by a single Cis-element and its initial functional characterization in vessels. Journal of molecular and cellular cardiology 138: 147–157.

48. Zhao, J, Wu, W, Zhang, W, Lu, YW, Tou, E, Ye, J, et al. (2017). Selective expression of TSPAN2 in vascular smooth muscle is independently regulated by TGF-beta1/SMAD and myocardin/serum response factor. Faseb j 31: 2576–2591.

49. Bryant, WB, Yang, A, Griffin, S, Zhang, W, Long, X, and Miano, JM (2022). CRISPR-LRS for mapping transgenes in the mouse genome. bioRxiv: 2022.2001.2005.475144.

50. Chen, J, Kitchen, CM, Streb, JW, and Miano, JM (2002). Myocardin: a component of a molecular switch for smooth muscle differentiation. Journal of molecular and cellular cardiology 34: 1345–1356.

51. Hon, CC, Ramilowski, JA, Harshbarger, J, Bertin, N, Rackham, OJ, Gough, J, et al. (2017). An atlas of human long non-coding RNAs with accurate 5’ ends. Nature 543: 199–204.

52. Tang, Y, Yang, X, Friesel, RE, Vary, CP, and Liaw, L (2011). Mechanisms of TGF-β-induced differentiation in human vascular smooth muscle cells. Journal of vascular research 48: 485–494.

53. Cai, Y, Nagel, DJ, Zhou, Q, Cygnar, KD, Zhao, H, Li, F, et al. (2015). Role of cAMP-phosphodiesterase 1C signaling in regulating growth factor receptor stability, vascular smooth muscle cell growth, migration, and neointimal hyperplasia. Circulation research 116: 1120–1132.

54. Ballantyne, MD, Pinel, K, Dakin, R, Vesey, AT, Diver, L, Mackenzie, R, et al. (2016). Smooth Muscle Enriched Long Noncoding RNA (SMILR) Regulates Cell Proliferation. Circulation 133: 2050–2065.

55. Harshe, RP, Xie, A, Vuerich, M, Frank, LA, Gromova, B, Zhang, H, et al. (2020). Endogenous antisense RNA curbs CD39 expression in Crohn’s disease. Nat Commun 11: 5894.

56. Kalota, A, Karabon, L, Swider, CR, Viazovkina, E, Elzagheid, M, Damha, MJ, et al. (2006). 2’-deoxy-2’-fluoro-beta-D-arabinonucleic acid (2’F-ANA) modified oligonucleotides (ON) effect highly efficient, and persistent, gene silencing. Nucleic Acids Res 34: 451–461.

57. Agostini, F, Zanzoni, A, Klus, P, Marchese, D, Cirillo, D, and Tartaglia, GG (2013). catRAPID omics: a web server for large-scale prediction of protein-RNA interactions. Bioinformatics (Oxford, England) 29: 2928–2930.

58. Olson, EN, and Nordheim, A (2010). Linking actin dynamics and gene transcription to drive cellular motile functions. Nature reviews Molecular cell biology 11: 353–365.

59. Fang, F, Yang, Y, Yuan, Z, Gao, Y, Zhou, J, Chen, Q, et al. (2011). Myocardin-related transcription factor A mediates OxLDL-induced endothelial injury. Circulation research 108: 797–807.

60. Senft, D, Qi, J, and Ronai, ZA (2018). Ubiquitin ligases in oncogenic transformation and cancer therapy. Nature reviews Cancer 18: 69–88.

61. Bhattacharya, U, Neizer-Ashun, F, Mukherjee, P, and Bhattacharya, R (2020). When the chains do not break: the role of USP10 in physiology and pathology. Cell death & disease 11: 1033.

62. Leung, I, Dekel, A, Shifman, JM, and Sidhu, SS (2016). Saturation scanning of ubiquitin variants reveals a common hot spot for binding to USP2 and USP21. Proceedings of the National Academy of Sciences of the United States of America 113: 8705–8710.

63. Wang, L, Wu, D, and Xu, Z (2019). USP10 protects against cerebral ischemia injury by suppressing inflammation and apoptosis through the inhibition of TAK1 signaling. Biochem Biophys Res Commun 516: 1272–1278.

64. Luo, P, Qin, C, Zhu, L, Fang, C, Zhang, Y, Zhang, H, et al. (2018). Ubiquitin-Specific Peptidase 10 (USP10) Inhibits Hepatic Steatosis, Insulin Resistance, and Inflammation Through Sirt6. Hepatology (Baltimore, Md) 68: 1786–1803.

65. Zhang, C, Zhao, H, Cai, Y, Xiong, J, Mohan, A, Lou, D, et al. (2021). Cyclic nucleotide phosphodiesterase 1C contributes to abdominal aortic aneurysm. Proceedings of the National Academy of Sciences of the United States of America 118.

66. Goodwin, LO, Splinter, E, Davis, TL, Urban, R, He, H, Braun, RE, et al. (2019). Large-scale discovery of mouse transgenic integration sites reveals frequent structural variation and insertional mutagenesis. Genome research 29: 494–505.

67. Fanucchi, S, Fok, ET, Dalla, E, Shibayama, Y, Borner, K, Chang, EY, et al. (2019). Immune genes are primed for robust transcription by proximal long noncoding RNAs located in nuclear compartments. Nat Genet 51: 138–150.

68. Liu, B, Sun, L, Liu, Q, Gong, C, Yao, Y, Lv, X, et al. (2015). A cytoplasmic NF-κB interacting long noncoding RNA blocks IκB phosphorylation and suppresses breast cancer metastasis. Cancer Cell 27: 370–381.

69. Mineo, M, Lyons, SM, Zdioruk, M, von Spreckelsen, N, Ferrer-Luna, R, Ito, H, et al. (2020). Tumor Interferon Signaling Is Regulated by a lncRNA INCR1 Transcribed from the PD-L1 Locus. Molecular cell 78: 1207–1223.e1208.

70. Agarwal, S, Vierbuchen, T, Ghosh, S, Chan, J, Jiang, Z, Kandasamy, RK, et al. (2020). The long non-coding RNA LUCAT1 is a negative feedback regulator of interferon responses in humans. Nat Commun 11: 6348.

71. Caramori, G, Adcock, IM, Di Stefano, A, and Chung, KF (2014). Cytokine inhibition in the treatment of COPD. Int J Chron Obstruct Pulmon Dis 9: 397–412.

72. Ning, Y, and Lenz, HJ (2012). Targeting IL-8 in colorectal cancer. Expert Opin Ther Targets 16: 491–497.

73. Lane, BR, Lore, K, Bock, PJ, Andersson, J, Coffey, MJ, Strieter, RM, et al. (2001). Interleukin-8 stimulates human immunodeficiency virus type 1 replication and is a potential new target for antiretroviral therapy. Journal of virology 75: 8195–8202.

74. Stemmler, S, Arinir, U, Klein, W, Rohde, G, Hoffjan, S, Wirkus, N, et al. (2005). Association of interleukin-8 receptor alpha polymorphisms with chronic obstructive pulmonary disease and asthma. Genes Immun 6: 225–230.

75. Ha, H, Debnath, B, and Neamati, N (2017). Role of the CXCL8-CXCR1/2 Axis in Cancer and Inflammatory Diseases. Theranostics 7: 1543–1588.

76. Bilusic, M, Heery, CR, Collins, JM, Donahue, RN, Palena, C, Madan, RA, et al. (2019). Phase I trial of HuMax-IL8 (BMS-986253), an anti-IL-8 monoclonal antibody, in patients with metastatic or unresectable solid tumors. J Immunother Cancer 7: 240.

77. Hoffmann, E, Dittrich-Breiholz, O, Holtmann, H, and Kracht, M (2002). Multiple control of interleukin-8 gene expression. Journal of leukocyte biology 72: 847–855.

78. Bennett, M, Ulitsky, I, Alloza, I, Vandenbroeck, K, Miscianinov, V, Mahmoud, AD, et al. (2021). Novel Transcript Discovery Expands the Repertoire of Pathologically-Associated, Long Non-Coding RNAs in Vascular Smooth Muscle Cells. Int J Mol Sci 22.

79. Asfaha, S, Dubeykovskiy, AN, Tomita, H, Yang, X, Stokes, S, Shibata, W, et al. (2013). Mice that express human interleukin-8 have increased mobilization of immature myeloid cells, which exacerbates inflammation and accelerates colon carcinogenesis. Gastroenterology 144: 155–166.

80. Posern, G, and Treisman, R (2006). Actin’ together: serum response factor, its cofactors and the link to signal transduction. Trends Cell Biol 16: 588–596.

81. Cheng, X, Yang, Y, Fan, Z, Yu, L, Bai, H, Zhou, B, et al. (2015). MKL1 potentiates lung cancer cell migration and invasion by epigenetically activating MMP9 transcription. Oncogene 34: 5570–5581.

82. Xu, H, Wu, X, Qin, H, Tian, W, Chen, J, Sun, L, et al. (2015). Myocardin-Related Transcription Factor A Epigenetically Regulates Renal Fibrosis in Diabetic Nephropathy. Journal of the American Society of Nephrology: JASN 26: 1648–1660.

83. Warthi, G, Faulkner, JL, Doja, J, Ghanam, AR, Gao, P, Yang, AC, et al. (2022). Generation and Comparative Analysis of an Itga8-CreER (T2) Mouse with Preferential Activity in Vascular Smooth Muscle Cells. Nat Cardiovasc Res 1: 1084–1100.

## References

1. Ballantyne, MD, Pinel, K, Dakin, R, Vesey, AT, Diver, L, Mackenzie, R, et al. (2016). Smooth Muscle Enriched Long Noncoding RNA (SMILR) Regulates Cell Proliferation. Circulation 133: 2050–2065.

2. Warthi, G, Faulkner, JL, Doja, J, Ghanam, AR, Gao, P, Yang, AC, et al. (2022). Generation and Comparative Analysis of an Itga8-CreER (T2) Mouse with Preferential Activity in Vascular Smooth Muscle Cells. Nat Cardiovasc Res 1: 1084–1100.

