## Supplemental material for "*INKILN* is a novel long noncoding RNA promoting vascular smooth muscle inflammation via scaffolding MKL1 and USP10"

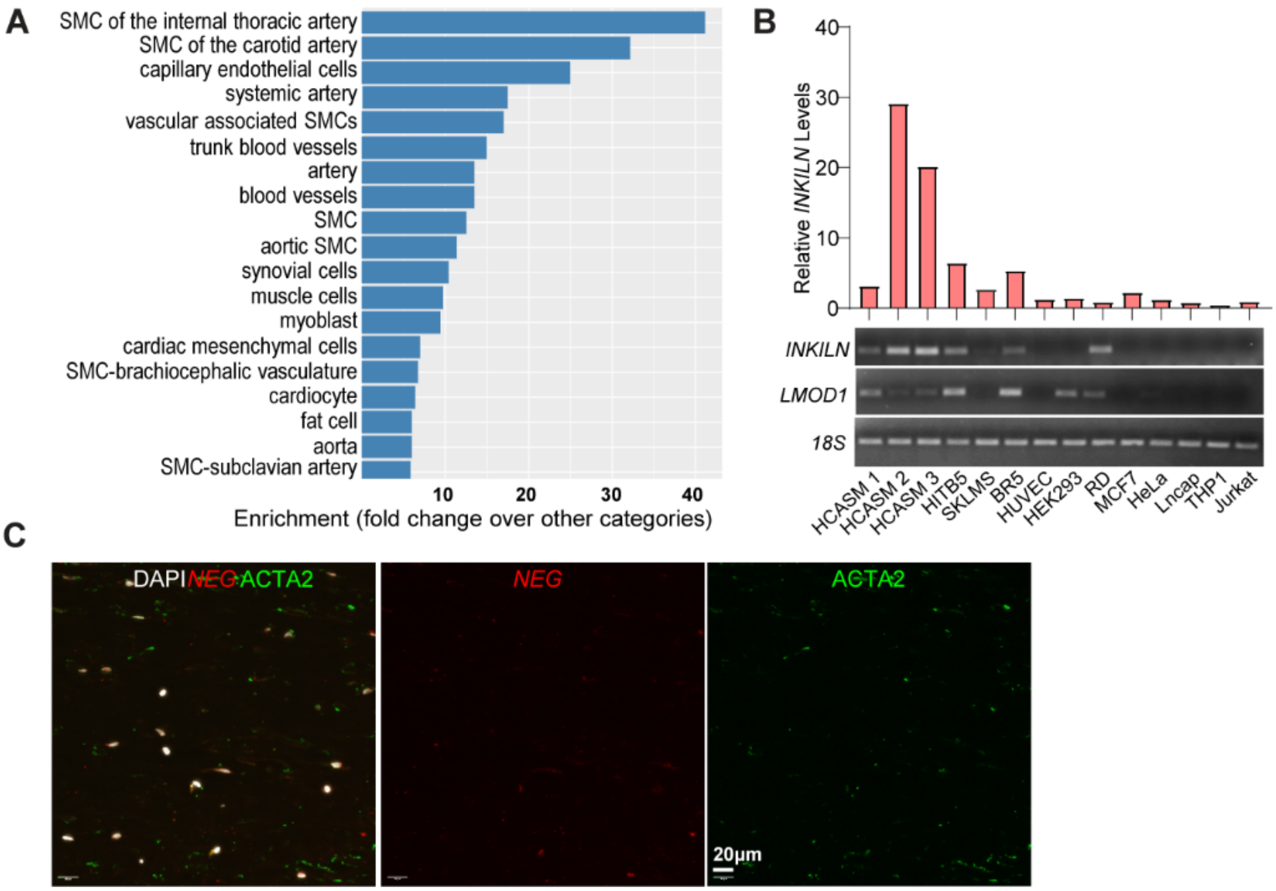

**Supplemental Figure 1. *INKILN* is enriched in cultured human SMCs and associated with the VSMC proinflammatory phenotype**. **A.** FANTOM (Functional Annotation Of the Mammalian genome) expression atlas analysis of 173 cell types and 174 tissues showed that *INKILN* was enriched in VSMCs and blood vessels. **B.** Semi-quantitative and quantitative RT-PCR (qRT-PCR) of the indicated genes in cultured human cells. **C.** Representative images of Immuno-RNA FISH for the negative control probe to *INKILN* and ACTA2 protein staining in the neointimal region of human abdominal aortic aneurysm (AAA) tissues (n=5 patients).

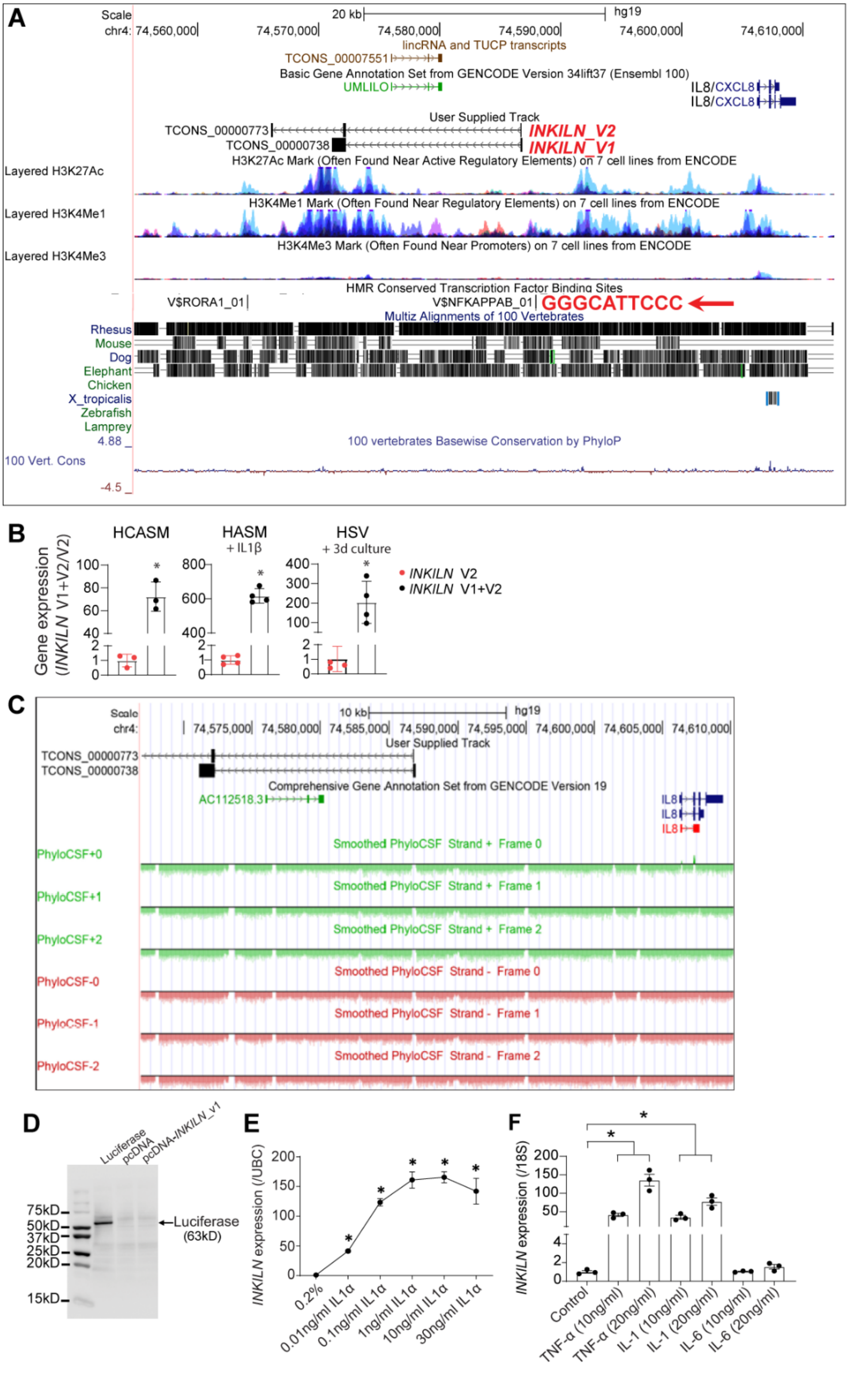

A
Scale
20 kb
hg19
chr4: 74,560,000 74,570,000 74,580,000 74,590,000 74,600,000 74,610,000
lincRNA and TUCP transcripts
TCONS_00007551
Basic Gene Annotation Set from GENCODE Version 34lift37 (Ensembl 100)
UMLILO
IL8/CXCL8
IL8/CXCL8
User Supplied Track
TCONS_00000773
INKILN_V2
TCONS_00000738
INKILN_V1
H3K27Ac Mark (Often Found Near Active Regulatory Elements) on 7 cell lines from ENCODE
Layered H3K27Ac
H3K4Me1 Mark (Often Found Near Regulatory Elements) on 7 cell lines from ENCODE
Layered H3K4Me1
H3K4Me3 Mark (Often Found Near Promoters) on 7 cell lines from ENCODE
Layered H3K4Me3
HMR Conserved Transcription Factor Binding Sites
V$RORA1_01
V$NFKAPPAB_01
GGGCATTCCC
Multiz Alignments of 100 Vertebrates
Rhesus
Mouse
Dog
Elephant
Chicken
X_tropicalis
Zebrafish
Lamprey
4.88
100 vertebrates Basewise Conservation by PhyloP
100 Vert. Cons
-4.5
B
HCASM
HASM
+ IL1β
HSV
+ 3d culture
Gene expression
(INKILN V1+V2/V2)
INKILN V2
INKILN V1+V2
C
Scale
10 kb
hg19
chr4: 74,575,000 74,580,000 74,585,000 74,590,000 74,595,000 74,600,000 74,605,000 74,610,000
User Supplied Track
TCONS_00000773
TCONS_00000738
Comprehensive Gene Annotation Set from GENCODE Version 19
AC112518.3
IL8
IL8
IL8
PhyloCSF+0
Smoothed PhyloCSF Strand + Frame 0
PhyloCSF+1
Smoothed PhyloCSF Strand + Frame 1
PhyloCSF+2
Smoothed PhyloCSF Strand + Frame 2
PhyloCSF-0
Smoothed PhyloCSF Strand - Frame 0
PhyloCSF-1
Smoothed PhyloCSF Strand - Frame 1
PhyloCSF-2
Smoothed PhyloCSF Strand - Frame 2
D
Luciferase
pcDNA
pcDNA-INKILN_v1
75kD
50kD
37kD
25kD
20kD
15kD
←Luciferase
(63kD)
E
INKILN expression (/UBC)
0.2%
0.01ng/ml IL1α
0.1ng/ml IL1α
1ng/ml IL1α
10ng/ml IL1α
30ng/ml IL1α
F
INKILN expression (/18S)
Control
TNF-α (10ng/ml)
TNF-α (20ng/ml)
IL-1 (10ng/ml)
IL-1 (20ng/ml)
IL-6 (10ng/ml)
IL-6 (20ng/ml)

**Supplemental Figure 2. *INKILN* noncoding potential validation and splicing variants characterization**. **A.** UCSC genome browser screen shot of the *INKILN* gene locus and its neighboring gene *CXCL8* (*IL8*). Two splice variants (*INKILN_V1* and *INKILN_V2)* were confirmed by sequence alignment and RACE analysis. The presence of active histone markers within the proximal 5'-end indicates that this region likely harbors an active promoter. Red arrow indicates a predicted conserved NF-κB site, which was validated in Figure 2J, 2K. **B.** qRT-PCR of *INKILN_V1+V2* versus *INKILN_V2* in the indicated human SMCs and ex vivo cultured human saphenous vein (HSV) segments (n=3). **C.** No predicted proteins/peptides were revealed within the *INKILN* gene locus by PhyloCSF. **D.** In vitro transcription/translation of pcDNA carrying *INKILN_V1*, pcDNA vector control, and a positive control luciferase expression plasmid and 2 μl of the protein mixtures were resolved in 16% Tricine SDS-PAGE gel and detected by Transcend Chemiluminescent Translation Detection system. An expected luciferase band (63kD) was seen in the positive control sample, but no translated product was seen in the sample derived from plasmid carrying *INKILN_V1*. Representative image is shown (n=3). **E.** qRT-PCR of dose-dependent induction of *INKILN* by IL1α in human saphenous vein SMCs (HSVSMCs) (n=3 biological replicates from one experiment). **F**. qRT-PCR of *INKILN* expression in pulmonary artery SMCs (PASMCs) under the indicated conditions for 24 hours (n=3 biological replicates from one experiment). **B**, unpaired two-tailed Student's t test; **F**, one-way ANOVA followed by multiple comparison test. $^{**}p<0.01$, $^{***}p<0.001$.

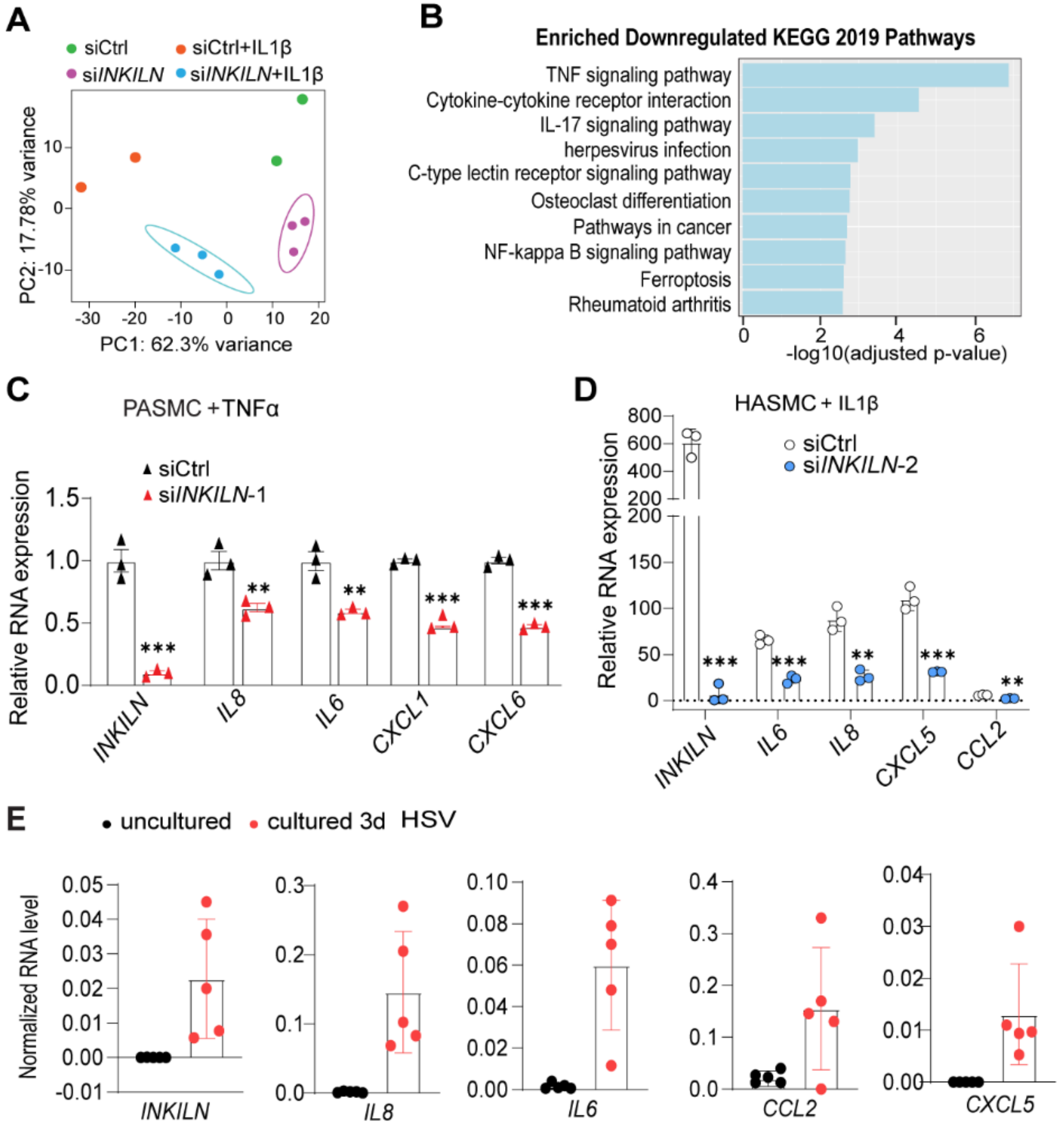

**Supplemental Figure 3. A.** Principal Component Analysis (PCA) of RNA-seq data shows distinctly clustered groups of HASMCs treated with 25 nM si*INKILN* or siCtrl (sicontrol) followed by IL1β (4 ng/ml) or vehicle control treatment for 24 hours. **B**. The top 10 KEGG pathways downregulated by si*INKILN-1* in HASMCs under the IL1β-induced condition are shown (adjusted p<0.05 and abs (log2FoldChange) from the bulk RNA-seq in **A**. **C**. qRT-PCR analysis of the indicated proinflammatory genes in PASMCs treated with si*INKILN-1* for 48 hours followed by TNFα (10 ng/ml) stimulation for 24 hours (n=3 technical replicates from 1 independent experiment). **D**. qRT-PCR analysis of the indicated proinflammatory genes in HASMCs treated by a second siRNA (si*INKILN*-2) for 48 hours followed by IL1β stimulation for 24 hours (n=3). Statistical analysis was done using an unpaired two-tailed Student's t test. ** p≤0.01, and

*** p≤0.001. **E.** qRT-PCR analysis of the indicated proinflammatory genes in uncultured versus ex vivo cultured HSV for 3 days (n=5).

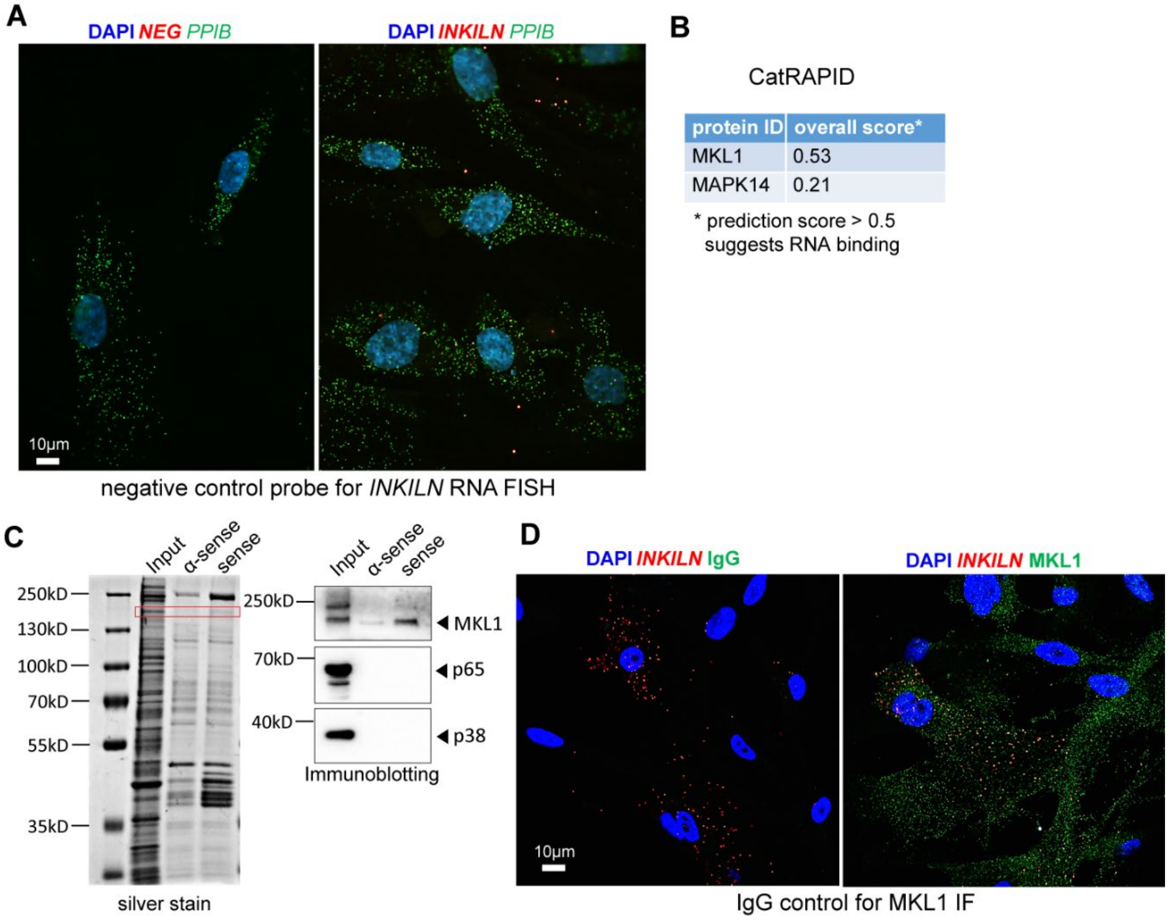

**Supplemental Figure 4. A.** RNA FISH using negative control probe to *INKILN* and positive control probe *PPIB* in HCASMCs. **B.** CatRAPID revealed MKL1 has greater RNA binding potential relative to MAPK14. **C.** Representative silver staining of the SDS-PAGE and western blot analysis for the indicated proteins from the precipitates of the in vitro transcribed antisense *INKILN* and sense *INKILN* incubated with protein lysates from HCASMCs (n=3). Red rectangle, highlighted bands with equivalent molecular weight of MKL1 protein. **D.** Negative control IgG for authentication of MKL1 immunostaining in HCASMCs.

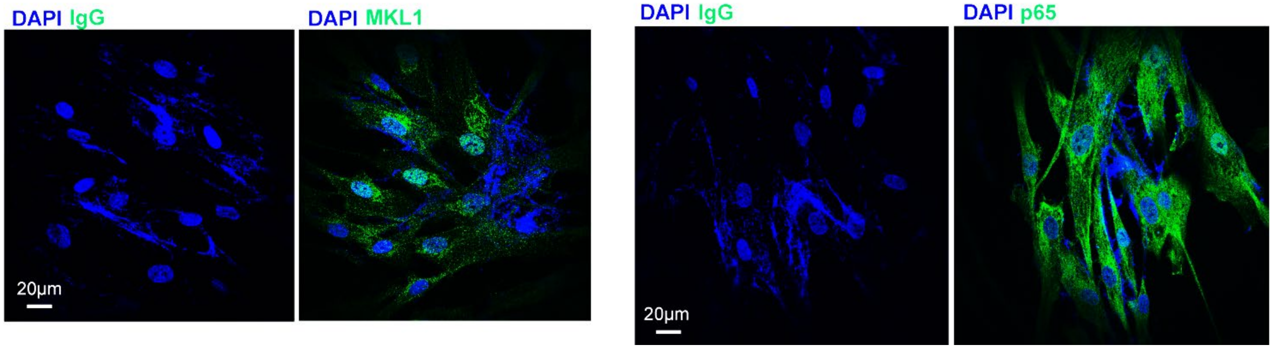

**Supplemental Figure 5.** Negative control IgG for authentication of MKL1 and p65 staining in HASMCs.

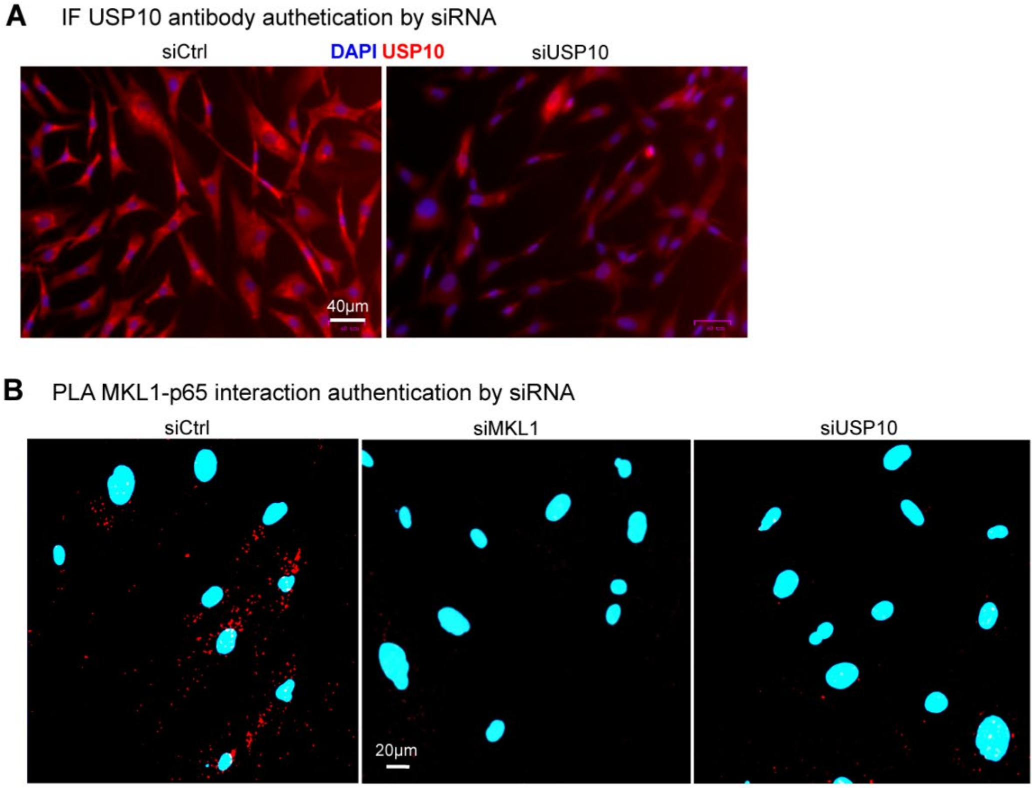

**Supplemental Figure 6**. **A.** Immunostaining for USP10 in HCASMCs ±si*USP10*. **B.** PLA was done in HCASMCs treated with siRNA control or siRNA to *MKL1* or *USP10*.

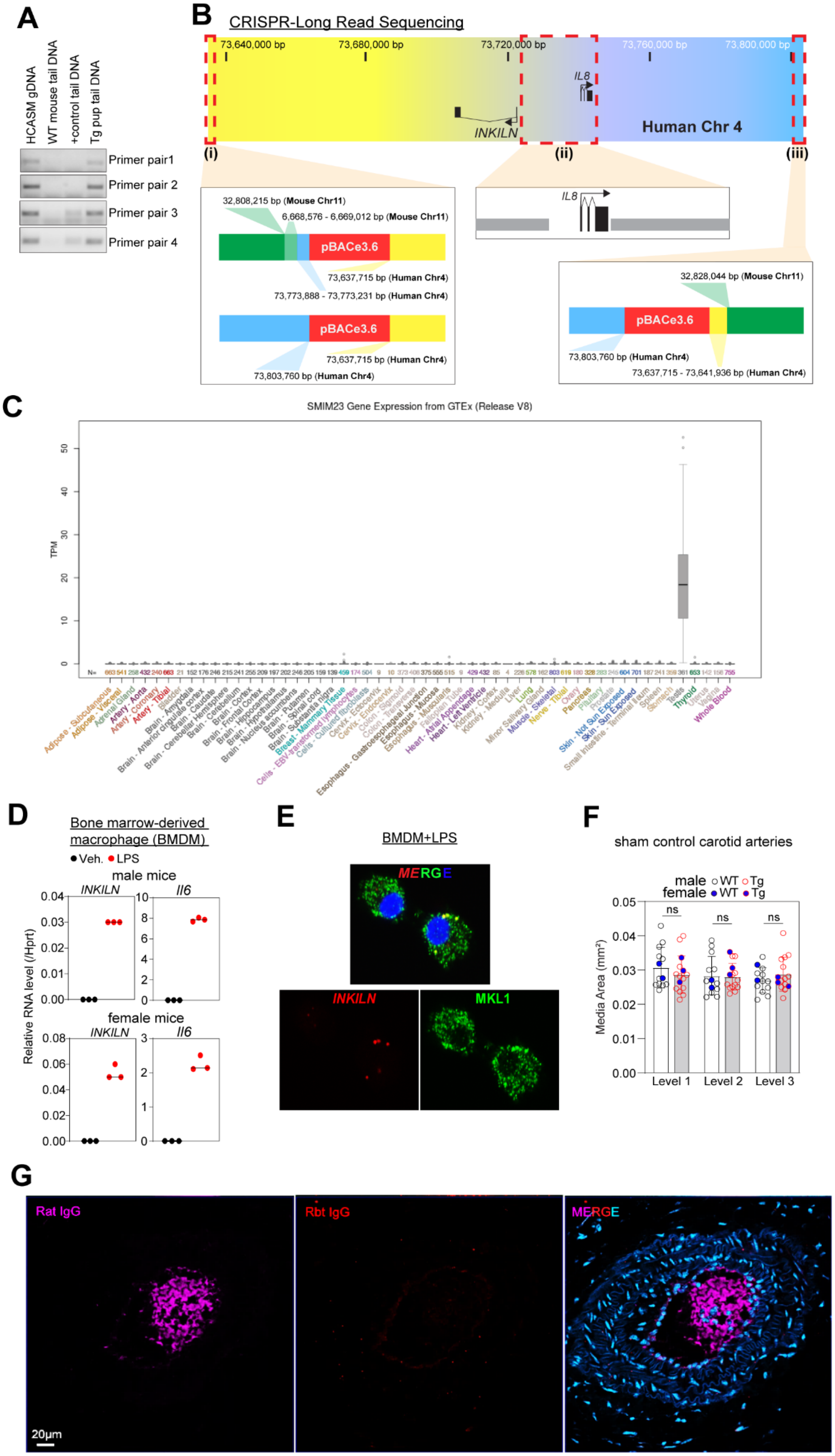
A
HCASM gDNA
WT mouse tail DNA
+control tail DNA
Tg pup tail DNA
Primer pair1
Primer pair 2
Primer pair 3
Primer pair 4
B
CRISPR-Long Read Sequencing
73,640,000 bp
73,680,000 bp
73,720,000 bp
73,760,000 bp
73,800,000 bp
IL8
INKILN
Human Chr 4
(i)
(ii)
(iii)
32,808,215 bp (Mouse Chr11)
6,668,576 - 6,669,012 bp (Mouse Chr11)
pBACe3.6
73,637,715 bp (Human Chr4)
73,773,888 - 73,773,231 bp (Human Chr4)
pBACe3.6
73,637,715 bp (Human Chr4)
73,803,760 bp (Human Chr4)
IL8
32,828,044 bp (Mouse Chr11)
pBACe3.6
73,803,760 bp (Human Chr4)
73,637,715 - 73,641,936 bp (Human Chr4)
C
SMIM23 Gene Expression from GTEx (Release V8)
TPM
Testis
Whole Blood
D
Bone marrow-derived macrophage (BMDM)
Veh. LPS
male mice
INKILN
Il6
female mice
INKILN
Il6
Relative RNA level (/Hprt)
E
BMDM+LPS
MERGE
INKILN
MKL1
F
sham control carotid arteries
male WT Tg
female WT Tg
ns
ns
ns
Media Area (mm²)
Level 1
Level 2
Level 3
G
Rat IgG
Rbt IgG
MERGE
20μm

**Supplemental Figure 7. A.** Genotyping the transgene *INKILN* using primers targeted to different regions of *INKILN* gene locus. **B.** RP11-997L11 BAC, which contains 73,637,715bp- 73,803,760bp of human chromosome 4 (hg38), was queried by CRISPR-LRS (i and iii) to map the *INKILN* transgene and (ii) to assess proper editing of the BAC. (i) Interrogating the 5' terminal end of the BAC defined one breakpoint of human and mouse genomes. CRISPR-LRS also found head to tail transgene tandem integration, indicating at least two copies. (iii) Interrogating the 3' terminal end of the BAC defined the other breakpoint of human and mouse genome sequences. (ii) Interrogating within the BAC demonstrated the complete removal of human *CXCL8* (*IL8*) and its promoter for proper study of human lncRNA *INKILN* in the mouse. **C.** GTEx data shows testis-specific gene expression of *SMIM23* in human tissues. **D.** qRT-PCR of *INKILN* and mouse *Il6* gene expression in bone marrow derived macrophages (BMDMs) from male and female *INKILN* Tg mice treated with LPS or vehicle control for 24 hours. **E.** Immuno-RNA FISH for MKL1 and *INKILN* in BMDMs from *INKILN* Tg mice induced by LPS for 24 hours. **F.** Quantitation of medial layer area at different levels in sham control carotids from WT versus *INKILN* Tg mice (n=12). Paired two-tailed Student's t test. ns, not significant. **G.** Species-matched negative control IgG for immunostaining of CD45 and MAC2; staining at center is non-specific.

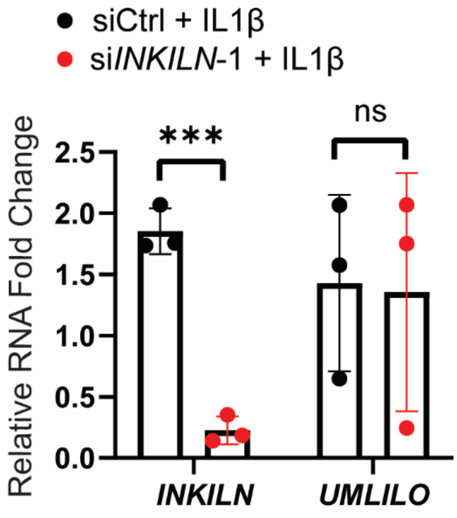

**Supplemental Figure 8. Loss of *INKILN* effect on *UMLILO* expression.** qRT-PCR of *UMLILO* in HASMCs transfected with si*INKILN*-1 versus siCtrl for 48 hours followed by IL1β treatment for 24 hours (n=3). Unpaired two-tailed Student's t test. ***p<0.001, ns: not significant.

**Supplementary Table 1. Patient characteristics for atherosclerosis from Munich Vascular Biobank**

| Characteristics | Atherosclerosis patients |
|---|---|
| Average age | 75.66 |
| Gender | 7 male /1 female |
| Hypertension (%) | 100% (8/8) |
| Diabetes (%) | 50% (4 diabetes/4 non-diabetics) |
| Anti-thrombotic therapy | 100% (8/8) |
| No active smokers | 5 out of 8 with smoking history (≥ 10 pack years) |

**Supplemental Table 2. Sequence information of oligos**

| Gene | Strand | Primer Sequence | Application(s) |
|---|---|---|---|
| *INKILN V1/V2* | Forward | ACCCTGTTTGTGAGGCTTC | qRT-PCR |
| | Reverse | TGCACTGGAAATGGCTGG | |
| *18S* | Forward | ATGGGCGGCGGAAAATAGC | qRT-PCR |
| | Reverse | TCTTGGTGAGGTCAATGTCTGC | |
| *CNN1* | Forward | ATGTCCTCTGCTCACTTCAAC | qRT-PCR |
| | Reverse | GCTGGTGGTCATACTTCTGG | |
| *LMOD1* | Forward | GCGGCAGAGAAACCAGAC | qRT-PCR |
| | Reverse | CCACTTGCTTGCTTTCATCC | |
| *MYH11* | Forward | ACGACAACTCCTCACGATTC | qRT-PCR |
| | Reverse | TCACTTCTCATCTTCTCCTTGG | |
| *IL8/CXCL8* | Forward | AGCCTTCCTGATTTCTGCAG | qRT-PCR |
| | Reverse | GTCCACTCTCAATCACTCTCAG | |
| *IL6* | Forward | GTGTTGCCTGCTGCCTTC | qRT-PCR |
| | Reverse | AGTGCCTCTTTGCTGCTTTC | |
| *CXCL1* | Forward | CTGCTCCTGCTCCTGGTAG | qRT-PCR |
| | Reverse | GCTTCCTCCTCCCTTCTGG | |
| *CCL2* | Forward | CTGTGCCTGCTGCTCATAG | qRT-PCR |
| | Reverse | CTTGCTGCTGGTGATTCTTC | |
| *CXCL5* | Forward | GCAAGTGTTCGCCATAGG | qRT-PCR |
| | Reverse | CGTGCTCATTTCTCTTAATCAG | |
| *IL1β* | Forward | AGGCACAAGGCACAACAG | qRT-PCR |
| | Reverse | GTGGTCGGAGATTCGTAGC | |
| *MYOSLID* | Forward | ACAGGGAGCCAGGACACC | qRT-PCR |
| | Reverse | GGAACCAGCACCAGGAACC | |
| *NEAT1* | Forward | GTGGTCTGAGGAGTGATGTG | qRT-PCR |
| | Reverse | GATAAGTTAAAGGGAGGAAGAAGG | |
| *RNU6-1* | Forward | GCT TCG GCA GCA CAT ATA CTA | qRT-PCR |
| | Reverse | CGA ATT TGC GTG TCA TCC TTG | |
| *IL1R1* | Forward | GATGAAGATGACCCAGTGCTAG | qRT-PCR |

| | | | |
|---|---|---|---|
| | Reverse | TGGCAAAACAGGTAAATGGATG | |
| *MKL1* | Forward | CAGCCTGAAGGAAGCCATC | qRT-PCR |
| | Reverse | GCCCATCGGAAGTTGAGAC | |
| *USP10* | Forward | TTTTAAATGCCACCGAACCTATC | qRT-PCR |
| | Reverse | CCAGCCATTCAGACCGATCT | |
| *UMLILO* | Forward | CTCAAACACTGGAGATAGAGCAG | qRT-PCR |
| | Reverse | TCACCGTGTACCAACTCTTTAC | |
| *INKILN* V1/V2 (636/637) | Forward | CCCTGTAAGATATGACTTGCTCC | qRT-PCR |
| | Reverse | TTCTGGTTCTGAAGATGTGGTC | |
| *INKILN* V2 | Forward | GTAAAGCAACATGCAGCCAG | qRT-PCR |
| | Reverse | AGAACCTGAAGCCAATGACA | |
| ENSG00000289530 | Forward | TCTAAAAGGATCACAGGCAGAAC | qRT-PCR |
| | Reverse | TGGTTCTGAAGATGTGGTCAC | |
| Smim23 | Forward | AGCAGAGACATGACAATCCAG | qRT-PCR |
| | Reverse | GTAGAGCACCAAGATGAGCAG | |
| *INKILN* Tg mice genotyping | Primer pair1-Forward | ACTGCAGACATCAGTCAGTG | PCR |
| | Primer pair1-Reverse | AGTGAATAATGACAGCTCATCCT | PCR |
| | Primer pair2-Forward | ATTGGATGGTTCACTTGTAGGA | PCR |
| | Primer pair2-Reverse | ACTGTCTATGTCGTGTCTTTAGG | PCR |
| | Primer pair3-Forward | GGATGCAGAGTTTACTGTGCA | PCR |
| | Primer pair4-Forward | CTTCAGGTTCTTCTTCCTCTGTC | PCR |
| | Primer pair3,4-Reverse | GTTTAAATAAAGTAGTCCAAGATCAT | PCR |
| DsNC1 | antisense | rCrArG rCrArA rUrUrA rGrCrG rCrArU rArUrU rArUrG rCrGrC rArUrA | Gene knockdown |
| | sense | rCrGrU rUrArA rUrCrG rCrGrU rArUrA rArUrA rCrGrC rGrUA T | |
| Dsi-*INKILN*-1 | antisense | rCrUrGrGrArArArUrGrGrCrUrGrGrCrUrGrCrArUrGrUrUrGrCrUrU | Gene knockdown |
| | sense | rGrCrArArCrArUrGrCrArGrCrCrArGrCrCrArUrUrUrCrCAG | |
| Dsi-*INKILN*-2 | antisense | rUrArC rArArG rUrGrU rArGrU rArArG rCrArU rGrUrU rArUrC rUrUrC | Gene knockdown |
| | sense | rArGrA rUrArA rCrArU rGrCrU rUrArC rUrArC rArCrU rUrGT A | |
| *INKILN* FANA ASO (5’-3’) | TGTAGTAAGCATGTTATCTTC | | Gene knockdown |
| *INKILN _V1 full length* | Forward | GATACGGATCCATTGAATTATGGGAGCGGGTATTTCCTGTGCTGT | PCR clone |
| | Reverse | GATACCTCGAGGGTCTGGTCTATGGATGATATTAGCTTATTTTAG | |
| *INKILN* -V2 full length | Forward | GAT ACG GAT CCCTCTCTCTT TGCCTGCCAC CCTGTAA | PCR clone |
| | Reverse | GAT ACC TCG AGTGAAGCATGTCTATAATATATTG | |

| | | | |
|---|---|---|---|
| *INKILN* -1.4 kb reporter | Forward | gatacggtaccTCAAAGTTCCCAGACCACTTC | PCR clone |
| | Reverse | gatacctcgagGCAAGGAGGAGCAAGTCATATC | |
| *INKILN* -1.18 kb reporter | Forward | gatacggtaccCCTAAAGACACGACATAGACAG | PCR clone |
| | Reverse | gatacctcgagGCAAGGAGGAGCAAGTCATATC | |
| *INKILN* ChIP NFkB site | Forward | TTTCAACAGGGGACAACTCC | qPCR |
| | Reverse | CTGTCTATGTCGTGTCTTTAGGAG | |

**Supplementary Table 3 Antibody information**

| **Primary Antibodies for Western Blot** | | | |
|---|---|---|---|
| **Target antigen** | **Vendor or Source** | **Catalog #** | **Working concentration** |
| MKL1 | Bethyl Laboratories | A302-201A | 1:1000 |
| MKL1/MRTF-A | Cell Signaling | 14760 | 1:1000 |
| GAPDH | Cell Signaling Technology | 5174 | 1:2000 |
| LMOD | Proteintech | 15117-1-AP | 1:500 |
| ACTB | Sigma-Aldrich | A5441 | 1:5000 |
| HA-Tag | Cell Signaling Technology | 3724S | 1:1000 |
| H3 | Abcam | Ab1791 | 1:1000 |
| TUBA | Sigma-Aldrich | T5168 | 1:5000 |
| Phospho-P65 | Cell Signaling Technology | 3033 | 1:500 |
| NF-κB p65 | Cell Signaling Technology | 8242 | 1:1000 |
| USP10 | Cell Signaling Technology | 8501 | 1:1000 |
| **Primary Antibodies for Immunofluorescence** | | | |
| **Target antigen** | **Vendor or Source** | **Catalog #** | **Working concentration** |
| MKL1 | Bethyl Laboratories | A302-201A | 1:1000 |
| MKL1 | Santa Cruz Biotechnology | Sc-390324 | 1:200 |
| USP10 | Cell Signaling Technology | 8501 | 1:200 |
| Galectin-3 (MAC2) | Thermo Fisher | 14-5301-82 | 1:200 |
| CD45 | BD Biosciences | 550539 | 1:50 |
| ACTA2 – Cy3 | Millipore Sigma | C6198 | 1:200 |
| Ki67 (D3B5) | Cell Signaling | 9129S | 1:30 |
| **Primary Antibodies for PLA** | | | |
| MKL1 | Bethyl Laboratories | A302-201A | 1:1000 |
| USP10 | Cell Signaling Technology | 8501 | 1:1000 |
| **Primary Antibodies for co-Immunoprecipitation** | | | |
| MKL1 | Bethyl Laboratories | A302-202A | 4ug/400uL lysate |
| Rabbit IgG ctrl | Abcam | Ab171870 | |
| **Primary Antibodies for RNA Immunoprecipitation** | | | |
| MKL1 | Bethyl Laboratories | A302-202A | 5ug/100uL lysate |
| USP10 | ThermoFisher Scientific | MA5-25766 | |
| IgG (included in Magna RIP kit) | Millopore Sigma | 17-700 | |
| **Secondary Antibodies for Western blot** | | | |
| Goat anti-Rabbit IgG (H+L) | Invitrogen/Thermo Fisher Scientific | 31460 | 1:5,000 |

| | | | |
|---|---|---|---|
| Secondary Antibody, HRP | | | |
| Rabbit anti-Mouse IgG (H+L), HRP | Invitrogen/Thermo Fisher Scientific | 31450 | 1:5,000 |
| Rabbit TrueBlot®: Anti-Rabbit IgG HRP | Rockland | 18-8816-31 | 1:500 |
| **Secondary Antibodies for Immunofluorescence** | | | |
| Alexa Fluor 488 goat anti-mouse IgG (H+L) | Invitrogen/Thermo Fisher Scientific | A11001 | 1:500 |
| Alexa Fluor 488 goat anti-rabbit IgG (H+L) | Invitrogen/Thermo Fisher Scientific | A11034 | 1:500 |
| Alexa Fluor 555 goat anti-rabbit IgG (H+L) | Invitrogen/Thermo Fisher Scientific | A27039 | 1:500 |

**Table 4 crRNA used to map BAC RP11-997L11**

| description | crRNA sequence minus PAM | associated library with SRA deposited data |
|---|---|---|
| targets 5' of RP11-997L11 | GTGTAGTAACCAAAATCGAA | *INKILN*_CRISPR-LRS_01 |
| targets *INKILN* edit | AGGTTCCCCCCATGACACGT | *INKILN*_CRISPR-LRS_01 |
| targets *INKILN* edit | GACGTACTCTTTGGTTACCT | *INKILN*_CRISPR-LRS_02 |
| targets *INKILN* edit | AACTCCAAACTCCACCGATT | *INKILN*_CRISPR-LRS_02 |
| targets *INKILN* edit | GTCATACTCCGTATTTGATA | *INKILN*_CRISPR-LRS_03 |
| targets 3' of RP11-997L11 | GCACAACCACCAAGTAAATG | *INKILN*_CRISPR-LRS_03 |
| targets 3' of RP11-997L11 | CTAAGTATCCTACTTCACCC | *INKILN*_CRISPR-LRS_04 |

**Table 5 Primers used for copy number of *INKILN* transgene analysis**

| description | sequence |
|---|---|
| 5'_internal-locus-control | CACCGGCTACACCAATCAA |
| 3'_internal-locus-control | CAGAAGATGTCTCCAGGCAAG |
| 5'_Cre-1 | CGTACTGACGGTGGGAGAAT |
| 3'_Cre-1 | CCCGGCAAAACAGGTAGTTA |
| 5'_*INKILN* | TGGTAGCAGGAGAGGAACTAATA |
| 3'_*INKILN* | CCTGCATTCACTCCCTTAACA |
